# Overcoming Genetic Drop-outs in Variants-based Lineage Tracing from Single-cell RNA Sequencing Data

**DOI:** 10.1101/2020.01.01.892562

**Authors:** Tianshi Lu, Seongoh Park, James Zhu, Xiaowei Zhan, Xinlei Wang, Hao Zhu, Tao Wang

## Abstract

Lineage tracing provides key insights into the fates of individual cells in complex tissues. Recent works on lineage reconstruction based on the single-cell expression data are suitable for short time frames while tracing lineage based on more stable genetic markers is needed for studies that span time scales over months or years. However, variant calling from the single-cell RNA sequencing (scRNA-Seq) data suffers from “genetic drop-outs”, including low coverage and allelic bias, which presents significant obstacles for lineage reconstruction. Prior studies focused only on mitochondrial (chrM) variants and need to be expanded to the whole genome to capture more variants with clearer physiological meaning. However, non-chrM variants suffer even more severe drop-outs than chrM variants, although drop-outs affect all variants. We developed strategies to overcome genetic drop-outs in scRNA-Seq-derived whole genomic variants for accurate lineage tracing, and we developed SClineger, a Bayesian Hierarchical model, to implement our approach. Our validation analyses on a series of sequencing protocols demonstrated the necessity of correction for genetic drop-outs and consideration of variants in the whole genome, and also showed the improvement that our approach provided. We showed that genetic-based lineage tracing is applicable for single-cell studies of both tumors and non-tumor tissues using our approach, and can reveal novel biological insights not afforded by expressional analyses. Interestingly, we showed that cells of various lineages grew under the spatial constraints of their respective organs during the developmental process. Overall, our work provides a powerful tool that can be applied to the large amounts of already existing scRNA-Seq data to construct the lineage histories of cells and derive new knowledge.

## INTRODUCTION

Lineage tracing provides a powerful means of understanding the cellular basis of development, regeneration, and disease^1–3^. Historically, tracing cell lineages in model organisms was achieved through low-throughput and invasive techniques such as labeling cells with dyes or radioactive tracers, or by the introduction of genetic elements^1^. Recent advances in single-cell sequencing have created opportunities to trace the lineage of a large number of cells and to follow their functional variations in a non-invasive manner^4^. More recently, spatial sequencing techniques, such as Slide-Seq^5^, were developed, which will further augment the power of lineage tracing by providing the spatial context while achieving near single-cell resolution.

In single-cell RNA-Seq (scRNA-Seq), the issue of drop-out in gene expression counting has been extensively studied^6,7^. Drop-out leads to a loss of signal when counting gene expression in scRNA-Seq data. Similarly, when scRNA-Seq or single-cell DNA sequencing (*e.g.* scATAC-Seq) are used for variant-based lineage tracing, “genetic drop-outs” become an important liability (**Sup. Fig. 1a**). There are two forms of “genetic drop-outs”: The first happens when the genotype of a genomic position is not detectable due to low coverage and/or a lack of expression. A variant may exist in all cells, but may only be detected in some. The second form of drop-out happens when only one allele in a cell is expressed or sampled. At the single-cell level, allele-specific expression is very common^8^. In addition, not all alleles may have been captured in the library due to randomness exacerbated by a low sequencing coverage. For a heterozygous variant in a cell, depending on which allele is expressed and/or sampled; this genomic position could be falsely identified as homozygous wild-type or homozygous mutated.

Recently, Ludwig *et al^9^* and Xu *et al^10^* demonstrated that mitochondrial (chrM) sequence variation could be used to infer clonal relationships from mito-sequencing and single-cell sequencing data. This is a clever approach because abundant, high depth chrM reads, which were previously ignored in downstream analyses, could now be exploited to trace lineages. However, there are significant limitations of this approach. Although the chance of detecting reliable variants in chrM is higher, the chromosomes are much larger compared to chrM. The focus on chrM variants ignores the opportunities to perform lineage tracing at a much finer resolution. Secondly, it is also difficult to correlate chrM variants with the physiological functions of the cells, but the impact of the non-chrM variants on the functions of the proteins and cells can be more readily assessed. In many research scenarios, such as tracking of mutations in tumor evolution, functional interpretability of variants is important. Therefore, it is critical to extend lineage tracing to non-chrM variants as well. However, although both chrM and non-chrM variants suffer from genetic drop-outs, the drop-outs affect non-chrM variants more than chrM variants (to be shown below).

Genetic drop-outs are a central issue that needs to be properly addressed before accurate lineages can be constructed from the single-cell-derived variants, especially non-chrM variants. We developed strategies to overcome genetic drop-outs in the scRNA-Seq-derived whole genomic variants, and developed SClineager, a Bayesian Hierarchical model, to implement this approach. We systematically validated our system on a suite of different sequencing protocols. We demonstrated the necessity of applying our approach for single-cell lineage tracing and its advantages over prior approaches. Our analyses showed that genetics-based single-cell lineage tracing, empowered by our system, is applicable for research studies involving both tumors and non-tumor tissues.

## RESULTS

### Detection of single-cell variants in the whole genome

There are inherent difficulties associated with the detection of variants in single cells. We first demonstrated the validity of the detected variants from the single cells and explored the nature of the non-chrM variants. We plotted the variant allele frequencies (VAFs, the number of variant reads divided by the number of total reads) for a total of 70 variants detected in the single cells of one chronic myeloid leukemia (CML) patient^11^ (Fig. 1a), with low coverage positions in cells shown as missing. In Fig. 1b, we showed the VAFs of three representative non-chrM variants across all cells, where peaks can be easily discerned, which correspond to homozygous wildtype, heterozygous mutant, and homozygous mutant genotypes.

**Fig. 1.**
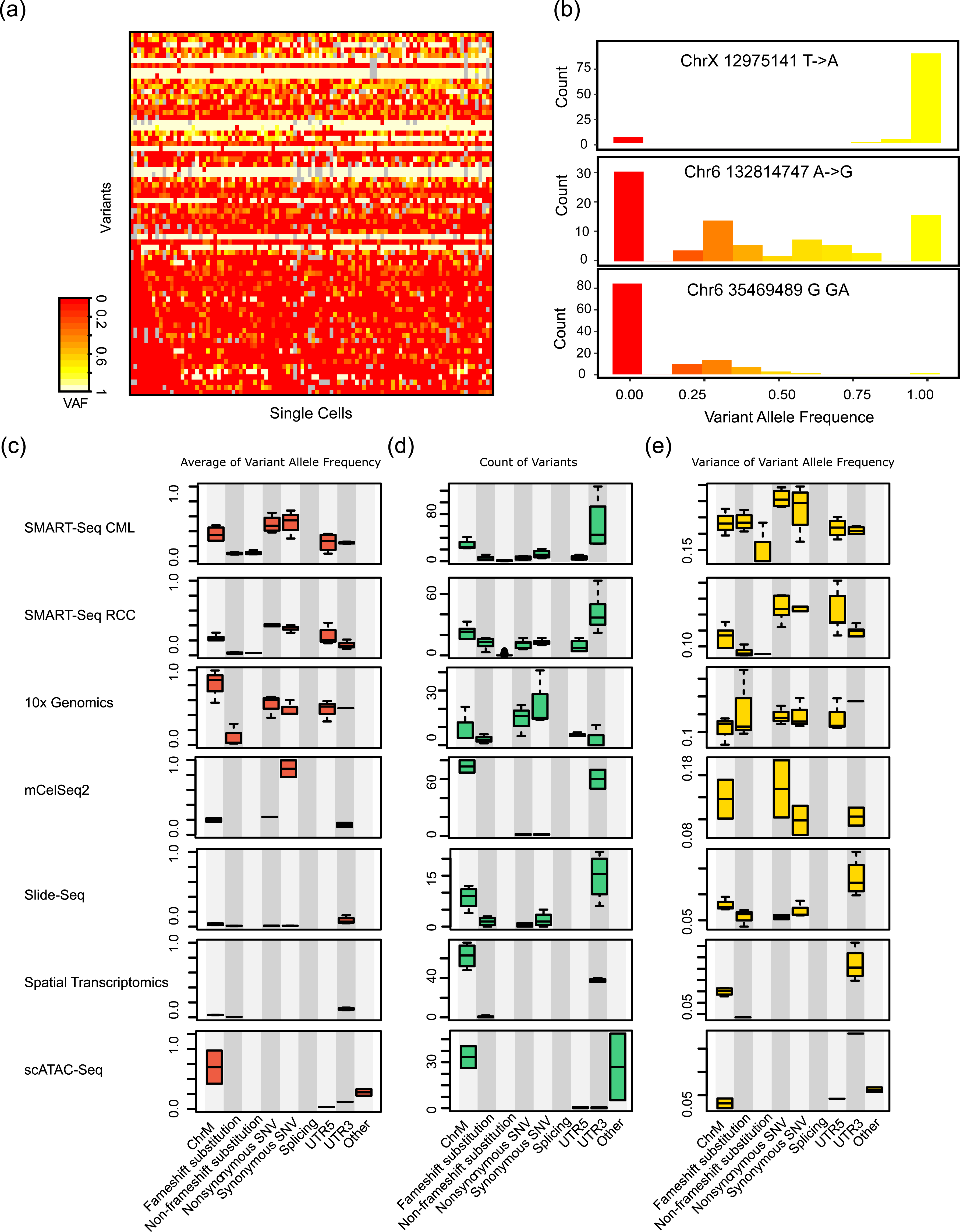
Reliable detection of single-cell variants in the whole genome. (a) Example Variant Allele Frequency (VAF) matrix of variants detected in all cells from one CML patient. Each row indicates one variant and each column indicates one single cell. Gray marks unobservable positions. Red means VAF=0 and yellow means VAF=1. The variants shown in the VAF matrix and used in the analyses of this study have already gone through the series of filtering steps that were outlined in **Materials and Methods**. (b) Example VAF histograms of several individual variants across all cells of (a). (c) Average VAF of each class of variants. (d) The number of each class of variants. (e) The variance of VAFs of each class of variants. The “Other” class refers to intronic and intergenic variants combined.

We analyzed datasets from a variety of sequencing protocols, including SMART-Seq, 10x Genomics, mCelSeq2, Slide-Seq, Spatial Transcriptomics (ST), and scATAC-Seq^5,9,11–15^. In Fig. 1c, we showed the VAFs of chrM variants, as well as variants on the chromosomes broken down by their functional classes. Variants of severe deleterious effects on transcripts (frameshift substitutions) led to low VAFs, potentially because the transcripts are more likely to be degraded. chrM, synonymous/non-synonymous single nucleotide variants (SNVs), and UTR variants have average VAFs of ~0.5. For the scATAC-Seq datasets, most detected non-chrM mutations were in intergenic and intronic regions (the “other” class), which is consistent with the whole genome sequencing manner of scATAC-Seq. Interestingly, variants of all classes from both Slide-Seq and Spatial Transcriptomics (ST) have relatively low VAFs. Slide-Seq and ST both perform sequencing on the beads to retain the physical locations of the sequences, which usually sample from a small group of cells, rather than exactly single cells. Therefore, only a portion of the cells in a given bead have a certain variant, which likely explains the low VAFs. Overall, these observations support the validity of our variant calling for these different sequencing techniques, which enables lineage tracing based on non-chrM variants in addition to chrM variants.

### Single-cell lineage tracing beyond mitochondrial variants

Leveraging the variants detected in the whole genome, rather than chrM variants alone, could have significant benefits for lineage tracing. We showed the number of each class of variants for each sequencing technique in Fig. 1d and **Sup. Fig. 2a**. We observed that the total number of variants on non-chrM chromosomes (all classes combined) were comparable or several folds more than the number of chrM variants, for all sequencing techniques investigated. This observation holds even after frameshift/non-frameshift substitutions, whose VAFs are subject to RNA degradation and thus are not optimal for lineage tracing, were removed (**Sup. Fig. 2a**).

Furthermore, we noticed (Fig. 1a) that many chrM variants’ VAFs were constant across all cells of the same sample, while non-chrM variants were more variable. To systematically evaluate this, we calculated the within-sample variance in VAFs of chrM variants and non-chrM variants for each sequencing technique (Fig. 1e). We observed that non-chrM variants (excluding substitutions) have comparable or much larger variability (depending on the type of variant and sequencing technique) than chrM variants. We calculated the proportion of the total genetic variability, represented by the sum of variances of all variants, explained by chrM and non-chrM variants for each sequencing technique, respectively (**Sup. Fig. 2b**). This practice shows that chrM variants only account for 0.08%-56.5% (average is 30.2%) of the total variance of all detected variants. These observations indicate that chrM variants may be better suited for cross-individual lineage tracing due to their stability with each individual, but for the problem of lineage tracing of cells within the same individual, non-chrM variants are more variable and can better mark different cell subsets.

### Genetic drop-outs in single-cell-derived chrM and non-chrM variants

Due to the technical challenges of the single-cell sequencing, the detected variants from single-cell sequencing suffer from genetic drop-outs. The first type of “genetic drop-out” is unobservability due to low coverage. In each single cell dataset, we counted the proportion of missing observations across all variants and all cells (Fig. 2a). It can be seen that 2%-20% of genomic positions (chrM and non-chrM) in the single cells are missing due to low coverage, depending on the sequencing technique. As expected, non-chrM variants have higher rate of missingness than chrM variants, despite that chrM variants also have drop-outs. Importantly, these analyses are already limited to variants of higher quality and better confidence from the series of filtering steps we employed (**Materials and Methods**). If we seek to avoid the issue of missingness by further narrowing down to variants that are completely observable across all cells, the sample sizes of variants that can be considered dramatically decrease (Fig. 2b). Therefore, a certain degree of missingness must be allowed to fully utilize the genetic information, but missingness needs to be carefully handled to correctly infer lineage.

**Fig. 2.**
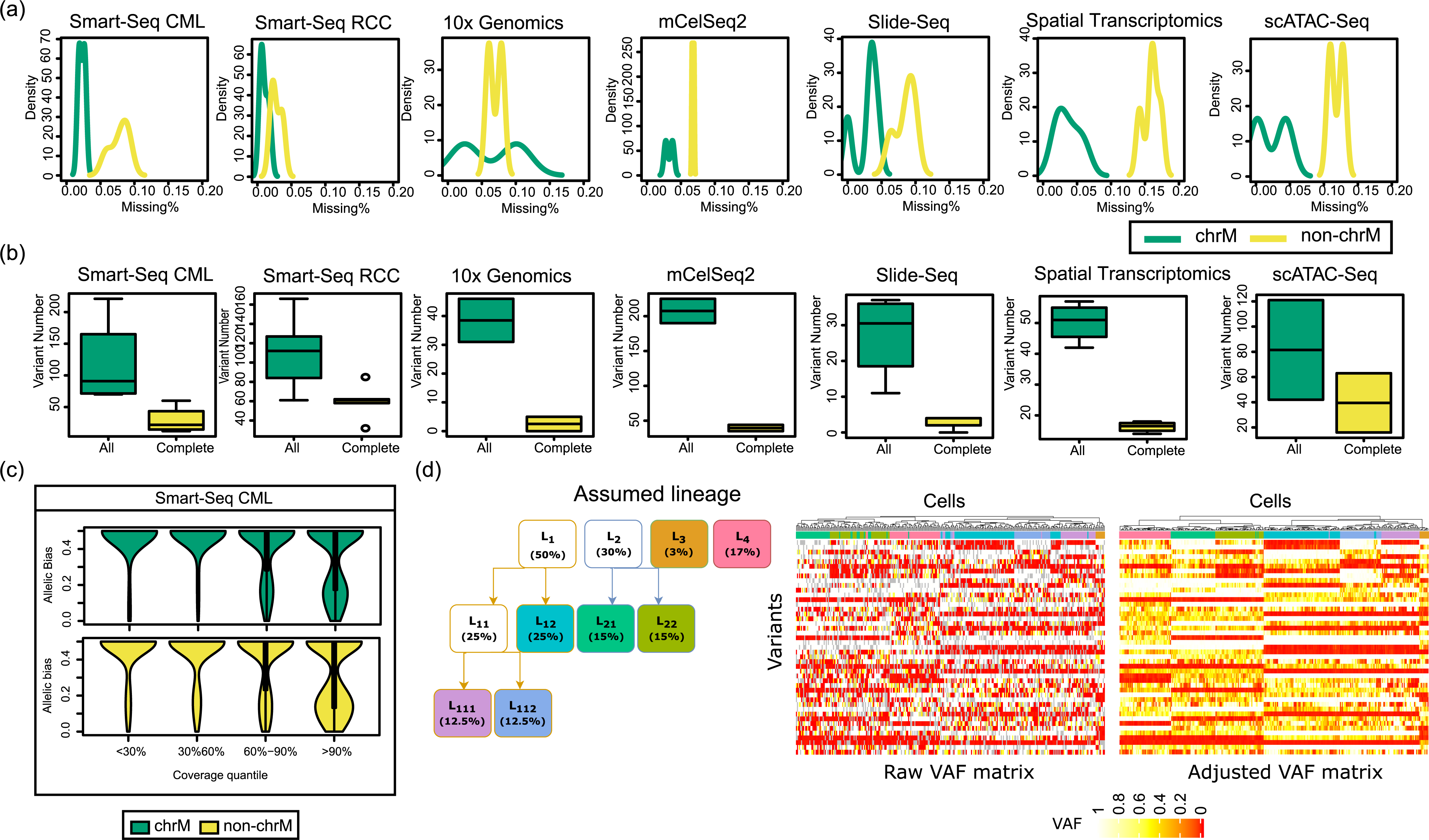
Handling genetic drop-outs in single cell variant calling and lineage tracing. (a) The first type of genetic drop-out – unobservability due to low coverage. The percentage of missing observations in the curated VAF matrix of each dataset was calculated, for chrM and non-chrM variants separately. (b) The number of all the variants that are in the VAF matrices and the number of variants that are completely observable across all cells, for each sequencing technique. (c) The second type of genetic drop-out – allelic imbalance. The X axis dichotomized the variants into bins of quantiles according to the coverage depths. The Y axis shows the distance between the VAF of each variant from 0.5. The results for the CML SMART-Seq dataset is shown. (d) Applying SClineager on a simulated dataset with 300 cells and 45 variants. The assumed true lineage structure is shown first. Then the raw VAF matrix and the SClineager-adjusted VAF matrix were shown as heatmaps. Each heatmap is ordered on the columns (cells) by unsupervised hierarchical clustering based on the VAFs of the variants. Missing positions are shown in gray.

Another form of “genetic drop-out” is wide-spread allelic imbalance in the single-cell sequencing data. For example, Borel *et al* observed that, in human primary fibroblasts, 76.4% of the heterozygous single nucleotide variants (SNVs) displayed stochastic monoallelic expression in single cells^16^. As it is difficult to determine the true genotype of each variant for each cell, we leveraged the correlation between expression level (sequencing depth) and the probability of allelic bias, an observation made by Fan *et al^17^*, to demonstrate allelic bias in the single-cell variants. Fan *et al* observed that more highly expressed genes in single cells are less likely to show allelic bias. The actual reason for this observation could be two-fold. First, it is likely true that highly expressed genes express both alleles (diploid organism for example), and *vice versa*. Another possibility is that in the library building process for sequencing, only transcripts of one genotype happen to be sampled and sequenced. This is more likely to happen to genes of lower expression and/or genomic regions of poorer coverage. The former effect applies to only scRNA-Seq data, while the second applies to scRNA-Seq and scATAC-Seq. Nevertheless, both effects should lead to a smaller probability of observing allelic imbalance when the sequencing count is higher. We calculated how much the VAF deviates from 0.5 for all variants of all datasets, and we showed these differences as a function of the sequencing coverage (Fig. 2c and **Sup. Fig. 3**). Overall, higher coverage is indeed associated with smaller differences between VAFs and 0.5 for the datasets investigated, confirming the significant impact of allelic bias on the single-cell variant detection. Importantly, this effect exists and is comparable for both chrM and non-chrM variants.

### Overcoming genetic drop-outs in single-cell sequencing for lineage tracing

To trace the lineage of single cells based on whole-genome variants while correcting for the two types of genetic drop-outs, we designed a Bayesian Hierarchical model, named SClineager (**Sup. Fig. 1b**). SClineager builds a hierarchical dependent relationship from the latent cell lineages to be inferred, to the variants with true VAFs to be inferred, and to the variants and their VAFs that are observed. In the observed data, some variants are lost due to low coverage, and the VAFs of the rest observable variants are subject to potential allelic imbalance. The core rationale of SClineager is that cells that are closer to each other on the genetic trajectory should have similar variants and VAFs, but the true variants/VAFs are not directly observed, due to missingness or allelic bias. However, the VAFs of the observed portion of the variants should only deviate from the true VAFs to some extent, and also in a patterned manner. SClineager considers two rules to capture the patterns: (1) variants with higher sequencing coverage should have less allelic imbalance, as is shown above; and (2) heterozygous variants are less likely to have allelic imbalance, as both alleles are already observed. In Fig. 2d, we applied SClineager on a simulated dataset where several lineages/sub-lineages of cells were assumed and the cells within the same lineage/sublineage have closely related genotypes. Without SClineager, unsupervised hierarchical clustering cannot correctly detect the different lineages of cells, while with SClineager, the clustering can correctly identify the cellular lineages with almost 100% accuracy. In addition, the genetic drop-outs have been corrected, and now the cells in the same lineage have very similar variants/VAFs. The mathematical description of SClineager is shown in **Sup. File 1**, along with more detailed simulation and convergence analyses.

We validated SClineager on the 969 CD45^+^ cells from the tumor microenvironment of 5 different Renal Cell Carcinoma (RCC) patients, which we processed by SMART-Seq. Patient origins served as the gold-standard labels in these cells. We showed a heatmap of SClineager-inferred VAFs for all 969 cells (Fig. 3a). It is evident that cells from different patients possess very unique sets of variants. We applied unsupervised hierarchical clustering of the VAFs of all variants and observed that cells from different patients clustered into tight groups (Fig. 3b). In comparison, we carried out the same analyses using the raw observed VAFs of all the variants, where missing positions were replaced by population medians, or using the raw VAFs of only the variants that are observable across all the cells. The same analysis showed that the clustering of cells is much less clear (Fig. 3c). We also carried out the analyses using SClineager-inferred VAFs, but using only chrM variants or only non-chrM variants (Fig. 3d). In both cases, we observed less clear clustering of cells by patient origins.

**Fig. 3.**
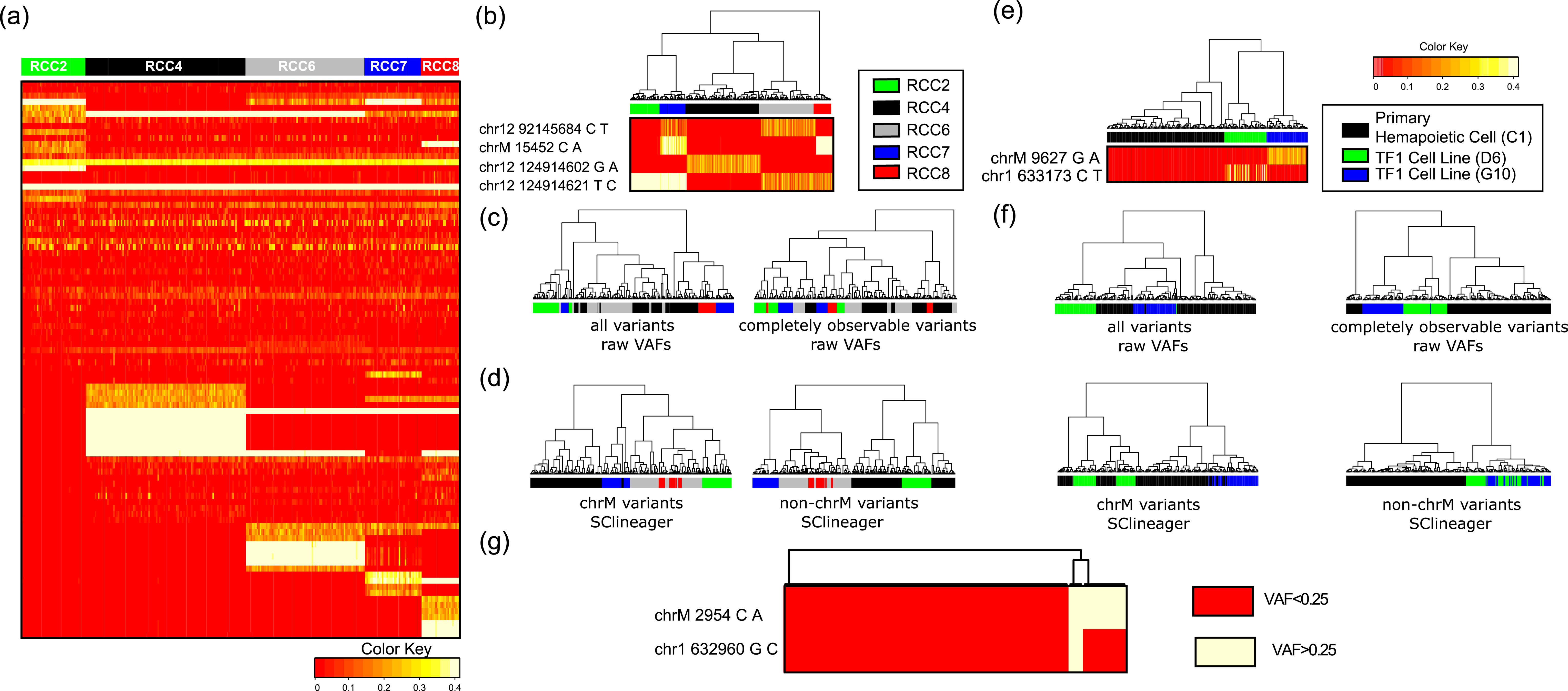
Validation of the SClineager algorithm. (a) A heatmap of SClineager-inferred variants and their VAFs for all 969 cells from the 5 RCC samples. (b) Unsupervised hierarchical clustering of SClineager-inferred VAFs of all variants shows the clear clustering of cells by their patient origin. (c) Hierarchical clustering analyses of the raw VAFs of all the variants (missing positions were replaced by population medians) or of the raw VAFs of only the variants that are observable across all the cells. (d) Hierarchical clustering analyses of the SClineager-inferred VAFs, but using only chrM variants or only non-chrM variants. (e) Unsupervised hierarchical clustering of SClineager-inferred VAFs of all variants shows the clear clustering of C1, D6 and G10 hematopoietic cells for the scATAC-seq dataset. (f) Comparison analyses for (e). The same comparison analyses as in (c) and (d) were carried out, and were shown in the same order. (g) Non-chrM variants can define lineages of cells with finer resolution. A heatmap of two variants (chrM 2954 C->A and chr1 632960 G->C) that further split the D6 clone into three sub-clones are shown.

We validated SClineager on a more complicated case of hematopoietic cell scATAC-Seq data from Ludwig *et al^9^*. These include 134 primary hematopoietic cells of one healthy individual and 95 cells of the TF1 cell line. The TF1 cell line has two clones, D6 (N=48) and G10 (N=47). Unsupervised hierarchical clustering of all the cells using SClineager-inferred VAFs of all variants show that the primary hematopoietic cells and the TF1 cells are very well separated (Fig. 3e, all variants shown in **Sup. Fig. 4**). More importantly, the two clones, D6 and G10, are also grouped in distinct sub-clusters of the TF1 cluster, suggesting that SClineager can distinguish lineages of cells from the same individual. We carried out similar comparison analyses (without SClineager correction, and with SClineager correction but considering only chrM and non-chrM, respectively) as above, and observed poor clustering of the primary hematopoietic cells and the two clones of the TF1 cell line (Fig. 3f). To further demonstrate the value of considering non-chrM variants, we focused on two variants (chrM 2954 C->A and chr1 632960 G->C) of the G10 clone (Fig. 3g). The chrM variant divided the G10 clone into two subsets, while the chr1 variant further dichotomized one of the clones into two smaller clones. Without the inclusion of the non-chrM variants, much less resolution can be achieved for dissecting the lineages of cells.

### SClineager detects the genetic evolution of single tumor cells

We applied SClineager on a series of datasets for further validation and for demonstrating its capability of deriving novel biological insights. We first focused on single tumor cells, as tumors undergo active mutagenesis, which provide sufficient materials for tracking lineages of cells based on mutations. We analyzed the chronic myeloid leukemia (CML) SMART-Seq dataset by Giustacchini *et al^11^*. We applied SClineager on the cells from the OX1931 patient and plotted a tSNE plot based on the variants of these cells (Fig. 4a). In this patient, the cells come from two batches, one before the blast crisis and another after the blast crisis. The two batches of cells are well mixed together on the genetic tSNE, suggesting that they are of the same cellular etiology. However, when we plotted these cells in the tSNE space of gene expression (Fig. 4a), pre-blast-crisis cells and blast-crisis cells are separated. In contrast, when cells from another patient (CML1266) were examined, we discovered a significant shift in the genetic background of the CML cells between the pre-blast and the blast stages (Fig. 4b), showing the existence of a different tumor evolutionary process for this patient compared with OX1931. We also examined individual variants. The cells from patient K562 form two clusters in the tSNE space of genetics (Fig. 4c). The variant, “chrM 5486 C->A”, mostly exists in one of the two clusters, and another variant, “chr17 50693598 A->G”, exists in the other cluster. Overall, SClineager uncovers novel knowledge regarding the genetic etiology of single tumor cells that is not afforded by investigation of only the expression of genes, and which could be informative for optimizing treatment decisions.

**Fig. 4.**
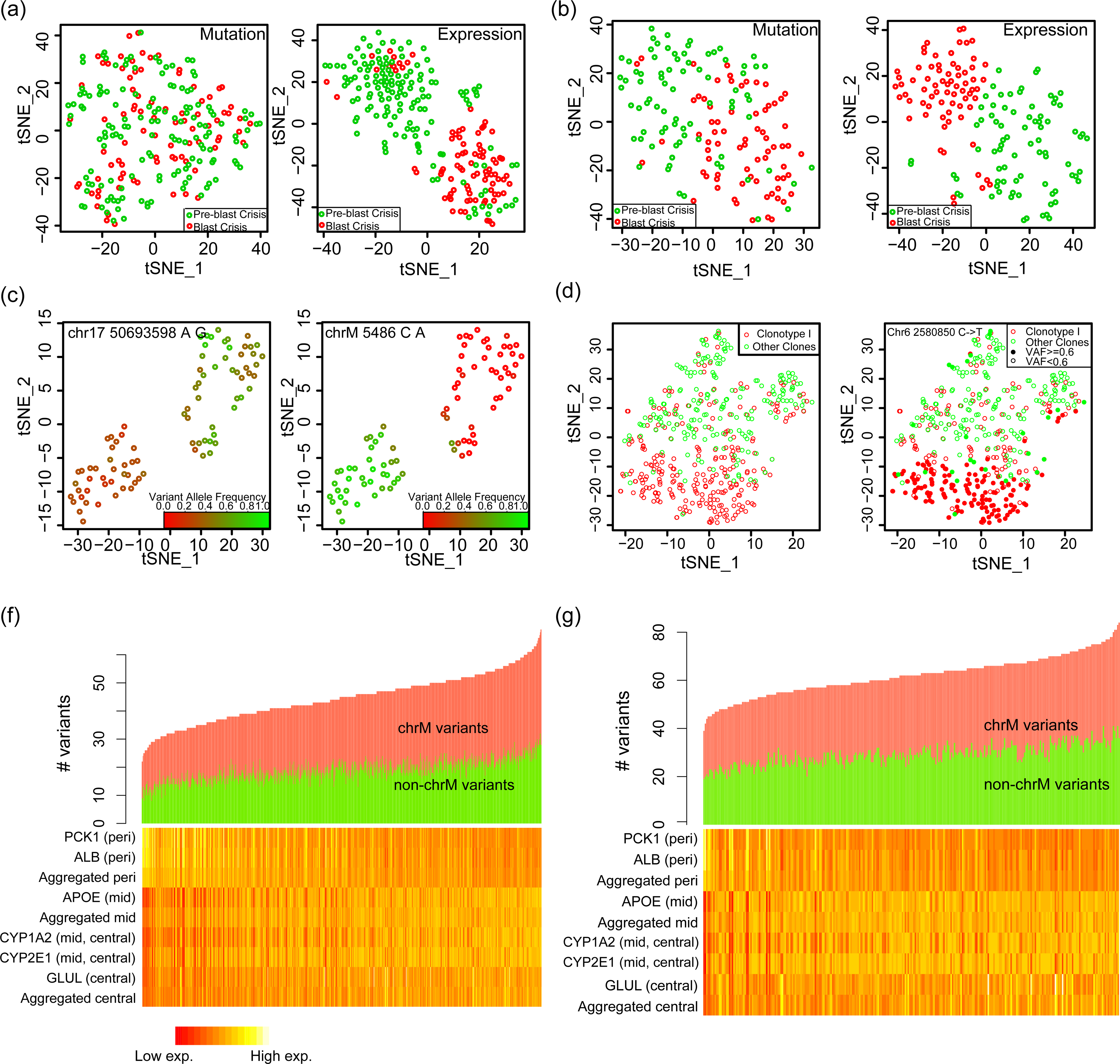
Discovery of novel biological insights through SClineager. (a) tSNE analysis of VAFs of the variants in the cells from the OX1931 CML patient and also the same analysis for gene expression. (a) tSNE analysis of VAFs of the variants in the cells from the CML1266 CML patient and also the same analysis for gene expression. (c) tSNE analysis of VAFs of all the variants in the cells from the K562 CML patient. The dots were colored according to the VAFs of the “chrM 5486 C->A” and “chr17 50693598 A->G” variants, respectively. (d) tSNE analysis of the VAFs of the mutations identified in the CTCL and non-malignant T cells. (e) The same tSNE plot as (d), but the dots are either solid or empty according to whether the specific variant is detected in that cell or not. (f) and (g) Waterfall plots showing the number of chrM and non-chrM mutations in liver hepatocytes and heatmaps showing expression of zonation landmark genes and average expression of zone-specific gene sets (“aggregated”). (f) P301 and (g) P304. The zone-specific gene sets are curated from gene modules defined by Aizarani *et al*. The central zone gene set is comprised of modules 31, 32, and 35; the mid-zone gene set is comprised of modules 22, 23, and 24; and the peri zone gene set is comprised of modules 1, 3, 13, and 14. We removed variants that correspond to common human SNPs and also variants that appeared in >70% of the sequenced cells, which are likely germline variants.

We applied SClineager on an ECCITE-seq dataset^13^. ECCITE-seq is a variant of the 10x single-cell sequencing technology that allows simultaneous detection of the transcriptome, proteins, and T cell receptor (TCR) clonotypes. Mimitou *et al* applied this technique on a cutaneous T-cell lymphoma (CTCL) patient. In CTCL, the tumor cells are T cells, sharing one TCR and originating from one founder T cell, and there are other non-malignant T cells. We analyzed this CTCL dataset using SClineager, and plotted the distribution of the T cells from this patient in the tSNE space of mutation VAFs. It is evident that the T cells with the most dominant TCR clone form one cluster, while the other T cells form another cluster (Fig. 4d), which indicates that the malignant T cells became genetically divergent from the indigenous T cell population after the tumorigenic event. We also identified variants that seem to be only found in the malignant T cell clone (*e.g.*, “chr6 2580850 C->T” shown in Fig. 4e). Interestingly, this mutation seems to mark two sub-clones of the CTCL cells, where one sub-clone is still genetically similar to the normal T cells, while the other sub-clone has become very divergent. We also carried out comparison analyses in this dataset to demonstrate the value of correction of genetic drop-outs. By the same analyses using the raw VAFs without applying SClineager (**Sup. Fig. 5**), we found that the malignant clone cannot be separated from the other clones. On the other hand, if we only examine the variants that are completely observable across all cells, we have only three mutations left, so tSNE cannot really be performed.

### SClineager identifies mutational hotspot zones in normal liver lobules

Next, we focused on single cells derived from non-malignant tissues, and demonstrated that non-chrM and chrM variants also accumulate during normal organ development or non-cancer disease processes that are amenable for our approach. We analyzed the human normal liver mCelSeq2 dataset by Aizarani *et al^14^*. The lobule is the histologic unit of the liver. Lobules are divided into three concentric zones spanning the portal to central vein axis: periportal, mid-lobular, and pericentral zones. In our prior work, we have shown that human livers are subject to constant generation of mutations even in the absence of liver cancers^18^. But it is not known which zone has the most active mutagenesis that could ultimately lead to liver cancer. Using SClineager, we examined the mutational rates in different zones of hepatocytes. Although this question in itself is not strictly lineage tracing, SClineager offers the benefit of a more accurate profiling of the variants and their VAFs from correction of genetic drop-outs.

We identified variants in the hepatocyte mCelSeq2 data and analyzed them *via* SClineager. In Fig. 4f (patient 301) and Fig. 4g (patient 304), we ranked the hepatocytes by their mutational load (after SClineager correction). It can be seen that cells with more chrM variants are also the cells with more non-chrM variants, confirming that we are investigating a genome-wide phenotype. We correlated, with the mutational loads, the expression of several landmark zonation genes as well as the average expression of zone-specific gene modules, defined by Aizarani *et al^14^* (Fig. 4fg and **Sup. Fig. 6**). Interestingly, the high expression of mid-zone markers and mid-zone gene modules correlated most positively with high mutational load, and it is followed closely by central-zone markers/modules and least by peri-zone markers/modules. These results indicate that the mid-zone of lobules undergoes the most active mutational process, which is more vulnerable to tumorigenesis. Very interestingly, Chen *et al^19^* recently discovered through orthogonal experimental approaches that the mid-zone hepatocytes are the main population of cells in the liver lobules that are undergoing active regeneration, which is also supported by Sun *et al^20^*. These discoveries are in alignment of our conclusion as the correlation between proliferation and mutagenesis has been well-established^21^.

### Single-cell lineages are spatially constrained by organ anatomy

We further validated the genetic lineages of normal cells by their spatial distribution, by using the ST and Slide-Seq datasets. The unit of sequencing in ST and Slide-Seq is the “bead”, which comprises a small population of cells. Although we cannot apply SClineager, which relies on a single cell assumption, the impact of both types of genetic drop-outs should be “auto-corrected” to some extent, as multiple cells are sequenced in each bead and the sequencing reads are aggregated. We analyzed the mouse olfactory bulb (MOB) ST data from Ståhl *et al^15^*. In this dataset, four neighboring coronal slices of the MOB of the same mouse were subjected to ST. **Sup. Fig. 7** shows a stained image of mouse MOB (Wikipedia) showing the glomerular layer (1^st^ layer), the external plexiform and mitral cell layer (2^nd^ layer), and the internal plexiform and granule cell layer (3^rd^ layer) of MOB. We called variants for each slice independently and showed that there was a large overlap of the variants from all four slices (**Sup. Fig. 8**). Strikingly, we discovered many variants whose spatial distributions strongly correlated with the anatomical structure of MOB. For example, the same “chrM 1899 G->A” variant is detected in 3 of all 4 slices. And across all these 3 slices, this variant only appeared in the 2^nd^ and 3^rd^ layers of MOB (Fig. 5a). On the other hand, we identified another variant “chr8 64592559 C->T”, which is located in the 1^st^ and 2^nd^ layers of MOB (**Sup. Fig. 9**). This indicates that the different layers of MOB are formed from cells of distinct genetic lineages.

**Fig. 5.**
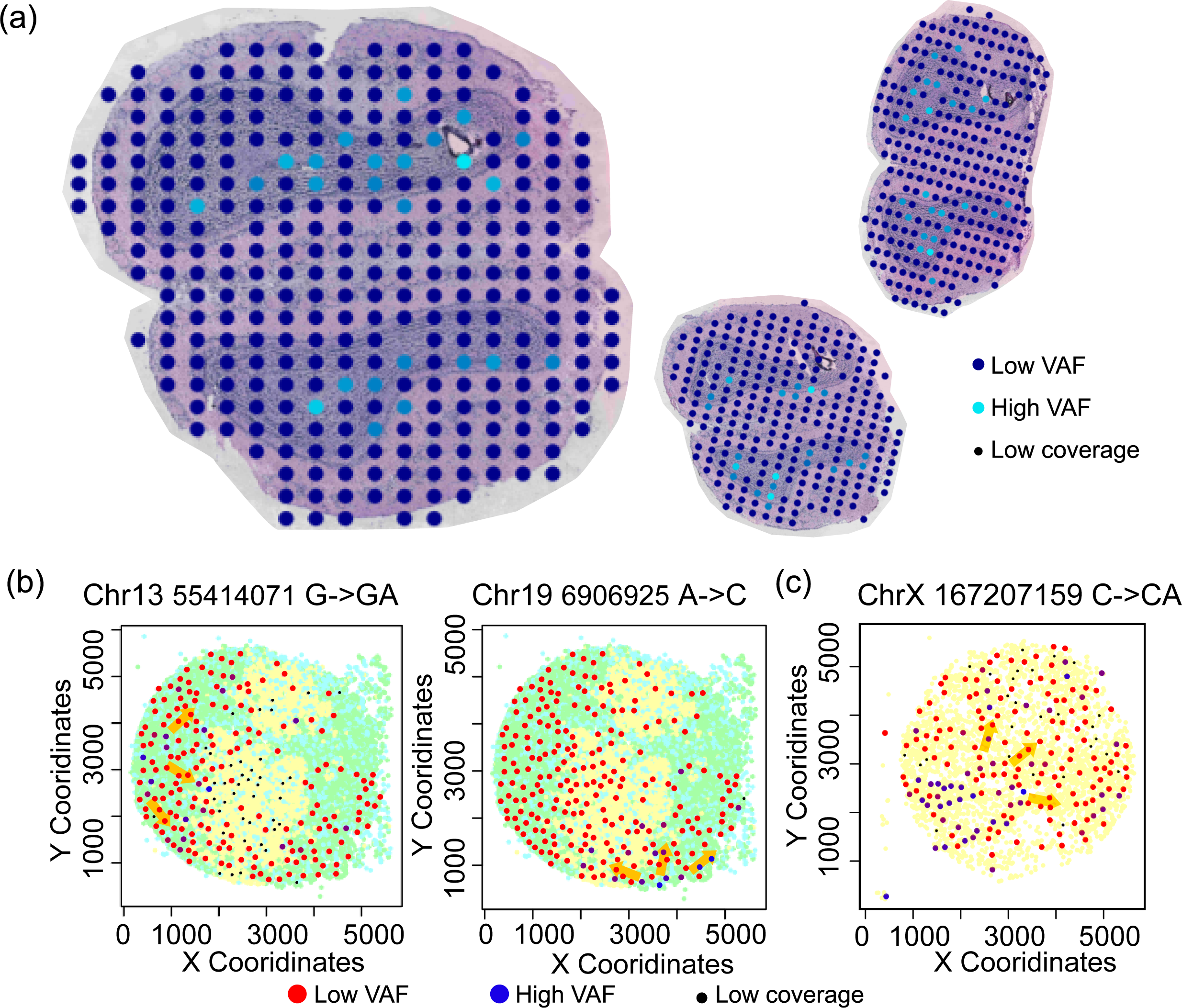
Cells of various lineages grew under the constraint of the organ anatomy. (a) Density and distribution of the “chrM 1899 G->A” variant in the three MOB slices where this variant is detected. Blue refers to high concentrations of this variant and cyan refers to low concentration of this variant. Small black dots refer to unobservable positions with insufficient coverage. (b) Density and distribution of the “chr13 55414071 G->GA” and “chr19 6906925 A->C” variants in the mouse kidney slide-seq dataset. Blue refers to high concentrations of this variant and red refers to low concentration of this variant. Small black dots refer to unobservable positions with insufficient coverage. (c) Density and distribution of the “chrX 167207159 C->CA” variant in the mouse hippocampus slide-seq dataset. For (b) and (c), the yellow arrows mark the direction of the VAF gradient. The unit of variant calling is “meta” beads that aggregated a number of neighboring beads (**Materials and Methods**). These original beads were also overlaid in (b) and (c) and were colored according to their group identities in unsupervised Kmeans clustering (**Sup. Fig. 9**), which likely represent different cell types.

Next, we analyzed the Slide-Seq data from Ludwig *et al^9^* for mouse kidney (Fig. 5b) and sagittal hippocampus (Fig. 5c). We observed a very intriguing spatial gradient pattern of the VAFs of the detected variants. Specifically, some variants appear in beads in a small region of the tissue slice with very high VAFs (reflecting high proportions of cells having a particular variant in one bead) and branch out to the beads in nearby regions with lower and lower VAFs. Fig. 5b shows two such variants, “chr13 55414071 G->GA” and “chr19 6906925 A->C”, in the mouse kidney, and Fig. 5c shows another variant, “chrX 167207159 C->CA”, in the mouse hippocampus. We performed unsupervised clustering based on gene expression (**Sup. Fig. 10**), and observed clear clusters of the beads from the mouse kidney, which represent different cell types. We overlaid the cluster identities of the beads upon the variant map for the mouse kidney, and observed that the distribution of the variants is correlated with the textural patterns of the kidney tissue, similar to our observation in Fig. 5a. For the hippocampus beads, we only detected one cluster (**Sup. Fig. 10**), and therefore didn’t perform this analysis. In the developmental process, one cell gains a variant and grows into a clone of cells. Our observation in mouse kidney and hippocampus suggests the existence of a physical “pushing and squeezing” effect of cells during development, which disseminates cells from the origins into nearby and distant regions, and finally forms a mosaic pattern of cells of various lineages. However, such an effect is not observed uniformly for all variants. We showed an example variant that is mostly homogeneously distributed among the beads (**Sup. Fig. 11**). Most such variants are detected in the majority of the beads, suggesting that they can be germline variants or very early somatic variants, which explains their wide and even distributions.

## DISCUSSION

We present the first study, to our knowledge, that formally characterizes genetic drop-outs in the single-cell variant calling. Strictly disregarding variants that are missing in only some of all the sequenced cells will lead to a severe loss of sample size. Neglect of allelic bias will lead to misleading construction of lineage trajectories. We presented solutions to address the genetic drop-out problems for lineage tracing. For most of the single-cell datasets, the SClingeager software we developed can handle both types of genetic drop-outs and perform lineage tracing based on genetic variants at the same time. For spatial sequencing protocols, like ST and Slide-Seq, genetic drop-outs are auto-corrected as their sequencing units are “beads”, which comprise a small group of cells so the sequencing coverage is increased and the stochastic allelic bias are averaged out. We systematically validated our approach of overcoming genetic drop-outs on a series of datasets, and demonstrated the significant advances that it provides.

Our work proved the necessity of considering variants in the whole genome, in addition to chrM, for lineage tracing. This will greatly expand the number of variants that can be considered, improve the interpretability of the physiological implication of the results, and is more suitable for intra-patient lineage tracing. The observation that non-chrM variants are more variable than chrM variants is probably due to the heteroplasmy of mitochondria^22,23^. There are usually hundreds of copies of chrM DNA in eukaryotic genomes, and it is common for mutations to affect only some copies. It is likely, in focused mito-Seq, these heteroplasmic variants can be readily detected and used for intra-patient lineage tracing of single cells. But according to our testing of commonly used single-cell sequencing protocols, it seems to be hard to detect such heteroplasmic variants due to their low VAFs. Therefore, it is critical to consider non-chrM variants for lineage tracing based on regular scRNA-seq data. However, genetic drop-outs must be carefully handled, which affect both chrM and non-chrM variants, but affect non-chrM variants more severely.

Lineage tracing can also be performed in the expressional space. A number of sophisticated and dedicated methods have been developed for this purpose^24–28^. However, these methods all operate under the rationale that the gradient changes in gene expression mark the cells’ locations on the trajectories. This assumption is true only in some scenarios that investigate biological phenomenon of shorter time scales, such as circadian rhythm. But many research questions require much longer tracking of cell lineages, over months or years (e.g. chronic disease follow-up). Methods for tracing of the single-cell lineages based on more stable genetic markers, are critically needed. Importantly, this work showed that analyses of variants and genetics-based lineage tracing of single cells is applicable not only for tumor tissues but also for non-tumor tissue types, ensuring the wide applicability of this approach.

However, mutagenesis in non-tumor tissues is admittedly much slower than the tumor tissues. A small number of mutations would hinder effective intra-sample lineage tracing, which relies on sufficient genetic differences between cells of the same sample. Therefore it’s even more important to include non-chrM variants in studies of normal tissues to increase the sample size of variants that can be considered. As we showed that genetic drop-outs are more striking in non-chrM variants compared with chrM variants, our strategies to overcome genetic drop-outs would provide critical benefit for lineage tracing studies of non-tumor tissues.

scRNA-Seq technologies have been developing rapidly over the past few years. But analytical methods for mining these data are still preliminary, and rich knowledge is left undiscovered in data archives and supplemental files of publications. Tracing the lineages of cells with genetic markers is one way of unleashing the full power of scRNA-seq. However, important issues like genetic-dropouts must be carefully dealt with for drawing informative conclusions. Our work has proposed solutions for genetic drop-outs, which should propel research and potentially translational applications involving lineage history construction from scRNA-Seq data.

## MATERIALS AND METHODS

### Splitting the raw sequencing files

The 10x Genomics single-cell sequencing platform outputs the raw sequencing reads of all cells in one Fastq file. We developed ScSplitter to split the reads in original Fastq file by their cell of origin using their cell barcode sequences. All reads were aligned to an appropriate reference genome using STAR^29^, and reads that fail to align were dropped. For reads sharing the same identical cell barcodes and UMIs, their alignment positions were compared and the reads that aligned to a position farther than the cutoff (default: 500 bp) from the median alignment position were removed. For paired reads, two median alignment positions were calculated and pairs that contain at least one read that fits the above criterion were kept. Output reads were grouped by cell identity and saved as separate Fastq files. At least 3,000 reads were required for a cell to be retained. The sequencing data for other platforms (ST, Spatial Transcriptomics, and mCelSeq2) were split in a similar manner.

### Variant calling algorithm

Variant calling was carried out for each cell independently. We used the QBRC mutation calling pipeline for variant calling in this study, developed in the Quantitative Biomedical Research Center (QBRC) of UT Southwestern Medical Center. In short, sequencing reads were aligned to the human or mouse reference genome by BWA-MEM^30^ for scATAC-seq and STAR^29^ for single-cell RNA sequencing. The STAR two-pass method was employed to handle read splitting caused by splicing of exons and introns. Picard was used to add read group information and sambamba was used to mark PCR duplicates. The variant calling was performed in the one-sample mode. Lofreq*^31^ was used for the ST and Slide-Seq datasets and Strelka2 Germline mode^32^ was used for other datasets. Annovar was used to annotate SNPs and Indels, and protein sequence changes^33^. All SNPs and Indels were combined and only kept if there were at least 5 total (wild type and variant) reads and at least 3 variant reads. The per-based coverage statistics were collected by “samtools mpileup”.

### Preprocessing of variant data

For datasets that are not from ST and Slide-Seq, sequencing data from each cell were subjected to variant calling. For ST, sequencing data from each bead were analyzed together. For Slide-Seq, each bead captured too few sequencing reads. Therefore, we pooled transcripts from neighboring beads to increase coverage and each “meta” bead is comprised of about 59.9 original beads and about 498k reads. For all datasets, variants detected in at least 3 cells (beads) and at least m% of all cells (beads), whichever is higher, are kept. m% is specifically determined for each sequencing technique, and ranges from 2%-10%. Then we removed variants whose variant allele frequencies (VAFs) are constant across all cells (beads). We tabulate all variants in a matrix where rows correspond to variants and columns correspond to cells (beads). For a given cell (bead) and a given variant, if the coverage in this cell (bead) at this genomic position is below a cutoff, c, we determined this position to be unobservable. c is specifically determined for each technique, and ranges from 3 to 20. For each row (variant), if this variant is unobserved in >a% of cells (beads), this variant is removed. For each column (cell or bead), if this cell (bead) has more than >b% of variants being unobservable, this cell (bead) is removed. a% and b% are determined specifically for each technique. The chosen values of all the cutoffs mentioned above for each technique/dataset of our study are shown in **Sup. Table 1**. In the end, we obtain a matrix of VAFs with rows being variants and columns being cells for each dataset. Unobservable positions are represented as missing values.

### SClineager

We developed a Bayesian hierarchical model to infer trajectories of single cells based on genetic markers. The essence of this model is to infer the true genotypes of the single cells, taking into account the fact that there are allelic bias and undetectable positions (the two types of drop-out), and to use the true genotypes to infer the lineage of cells. This inference task can be naturally handled by a Bayesian hierarchical model to incorporate the hierarchical relationship between (1) genetics-based correlation of the different cell lineages, (2) the true SNVs, which depend on the cell lineages, and (3) the observed SNVs, which depend on the true SNVs, but are subject to genetic drop-outs. The estimated correlation matrix for the cells will then be used to detect cell lineages by clustering methods. Our implementation of SClineager also includes two optional filters of variants before the model estimation starts. One filters out variants identified in specific genes. For example, the user may want to filter out genes located in low-complexity regions of the genome, which tends to cause problems for the read alignment and variant calling. The other filters out variants that correspond to common SNPs (population frequency for ESP6500 >0.01), to focus on variants that are more likely to be somatically acquired. A detailed description of the model setup is described in **Appendix I**.

### Statistical analyses

All computations and statistical analyses were carried out in the R computing environment. For all boxplots appearing in this study, box boundaries represent interquartile ranges, whiskers extend to the most extreme data point which is no more than 1.5 times the interquartile range, and the line in the middle of the box represents the median. The hierarchical clustering (for scATAC-Seq data) was performed using manhattan distance and the mcquitty clustering method. Single cell expression data (for the RCC SMART-Seq and mCelSeq2 datasets) was log(x+1) transformed. The SMART-Seq data were quantile normalized by the R *normalize.quantiles* function, while the mCelSeq2 data were normalized by library size. tSNE analyses were performed using the R *Rtsne* function. All P values were calculated in a two-way manner. The Slide-seq expression data were visualized on the tSNE plots by the R package *Rtsne.* The beads were clustered by R *kmeans* function.

### Single cell datasets used in this study

We generated the SMART-Seq datasets of 969 CD45^+^ cells from the tumor microenvironment of 5 kidney cancer (RCC) patients^12^. The SMART-Seq datasets of CML cells were created by Giustacchini *et al^11^*. We analyzed the top three patients with the most cells available, OX1931, CML1266, and K562. The 10X Genomics single cell sequencing datasets include the vdj_v1_hs_nsclc and the vdj_v1_hs_pbmc2 datasets from https://www.10xgenomics.com/resources/datasets/ and one ECCITE-seq dataset by Mimitou *et al^13^*, where ECCITE-Seq is a variant of the 10x single-cell sequencing technique. We analyzed the hepatocytes of two patients (P301 and P304), generated by Aizarani *et al via* the mCelSeq2 protocol^14^. These two patients have the majority of hepatocytes identified in this dataset. The definition of hepatocytes follows the definition of the original authors based on gene expression profiles. The Slide-Seq datasets of mouse kidney and brain tissues were extracted from the original Slide-Seq study^5^. Four pucks were analyzed, 180528_23 (kidney), 180611_7 (sagittal hippocampus), 180618_24 (sagittal hippocampus) and 180618_20 (sagittal hippocampus). The ST dataset of mouse olfactory bulbs was published by Ståhl *et al^15^*. We downloaded the raw data (MOB replicate 1-4) from https://www.spatialresearch.org/resources-published-datasets/doi-10-1126science-aaf2403/. We downloaded the raw scATAC-Seq datasets of primary hematopoietic cells and TF1 cells generated by Ludwig *et al^9^*.

## Code availability

QBRC mutation calling pipeline: https://github.com/tianshilu/QBRC-Somatic-Pipeline

The SClineager software: https://github.com/inmybrain/SClineager

Software for splitting the raw combined Fastq files: https://github.com/zzhu33/scSplitter

**Sup. Fig. 1** Genetic drop-out in single cell RNA sequencing data. (a) The top and bottom panels show the two types of genetic drop-outs. Yellow circles represent the nucleus with DNAs and grey circles represent cytosol with RNAs. “True” shows the true genotype type (DNA), and “Detected” shows the genotype inferred from scRNA-Seq data. (b) The model schema for SClineager.

**Sup. Fig. 2** chrM variants are insufficient for single cell lineage tracing. (a) The number of chrM and non-chrM variants detected for each sequencing technique. All chrM variants and non-chrM variants with or without indels excluded are shown. (b) Proportion of genetic information, represented by the proportion of variance of VAFs of all variants considered, accounted for by chrM and non-chrM variants, respectively.

**Sup. Fig. 3** The second type of genetic drop-out – allelic imbalance. The results for the RCC Smart-Seq, 10x Genomics, mCelSeq2, Slide-Seq, Spatial Transcriptomics, and scATAC-Seq datasets are shown. The format of the figure panels follow that of Fig. 2c.

**Sup. Fig. 4** Unsupervised hierarchical clustering of all scATAC-Seq C1, D6, and G10 cells using SClineager-inferred VAFs. All the variants are shown in the heatmap.

**Sup. Fig. 5** Comparison analyses for Fig. 4d. The left panel shows the tSNE plot of cells using the raw VAFs of variants (without SClineager adjustment), and right panel labels the existence of the “Chr6 2580850 C->T” mutation in the cells, which is investigated in Fig. 4d.

**Sup. Fig. 6** Scatterplots showing the mutational loads of hepatocytes and the expression of zonation landmark genes and average expression of zone-specific gene modules. The results for patient P301 are shown. The results for the total number of variants, the number of chrM variants, and the number of non-chrM variants were shown separately. The gene modules were defined in the same manner as in Fig. 4.

**Sup. Fig. 7** Anatomy of MOB. Image from https://en.wikipedia.org/wiki/Olfactory_bulb.

**Sup. Fig. 8** Venn diagram of the overlap of variants identified in the four slices of MOB ST datasets.

**Sup. Fig. 9** Density and distribution of the “chr8 64592559 C->T” variant in the four MOB slice ST data. Blue refers to high concentrations of this variant and cyan refers to low concentration of this variant. Small black dots refer to unobservable positions with insufficient coverage.

**Sup. Fig. 10** Unsupervised clustering of the Slide-seq datasets. Expression data of the original beads (pucks) from the two Slide-seq datasets that we investigated were plotted in the tSNE space. It is visually obvious that there are 3 clusters in the mouse kidney and only one cluster in the hippocampus. K-means clustering was employed to assign the cluster labels for the mouse kidney beads.

**Sup. Fig. 11** An example of homogeneously distributed variants identified in the mouse Slide-Seq data. Blue refers to high concentrations of this variant and red refers to low concentration of this variant. Small black dots refer to unobservable positions with insufficient coverage. The original beads were also overlaid as in Fig. 5 and were colored according to their group identities in unsupervised Kmeans clustering (**Sup. Fig. 9**), which likely represent different cell types.

**Sup. Table 1** Cutoffs employed for post-processing of variant data for each technique/dataset

**Sup. File 1** Detailed description of the statistical model setup of SClineager and simulation/diagnostic analyses

## ACKNOWLEDGEMENTS

We acknowledge Drs. Sergei Koralov, Anthony Cheng and Bo Yao for helping with the data transfer of the mCelSeq2 dataset involved in this study.

## AUTHOR CONTRIBUTIONS

T.L. carried out the primary data analyses. S.P. and J.Z contributed software packages. X.W., and H.Z. provided input on the study design. X.Z. downloaded the Slide-Seq data. T.W. supervised the whole study.

## COMPETING INTERESTS

The authors declare no conflicts of interest.

## Notes

**Funding:** This study was supported by the National Institutes of Health (NIH) [CCSG 5P30CA142543/TW], and Cancer Prevention Research Institute of Texas [CPRIT RP190208/TW]

