## Supplementary figures and images for "Overcoming Genetic Drop-outs in Variants-based Lineage Tracing from Single-cell RNA Sequencing Data"

### Supplemental Figure 1

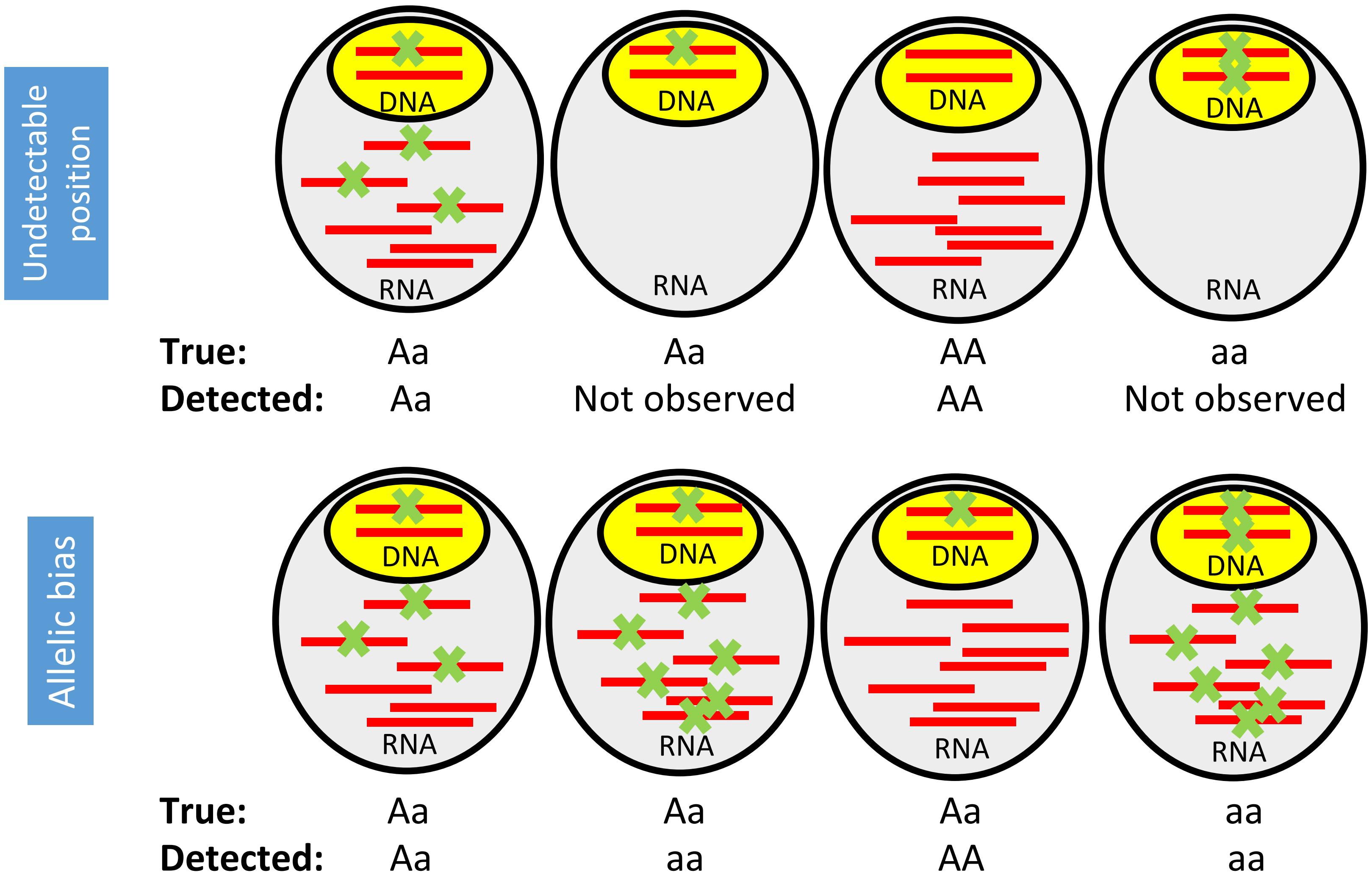

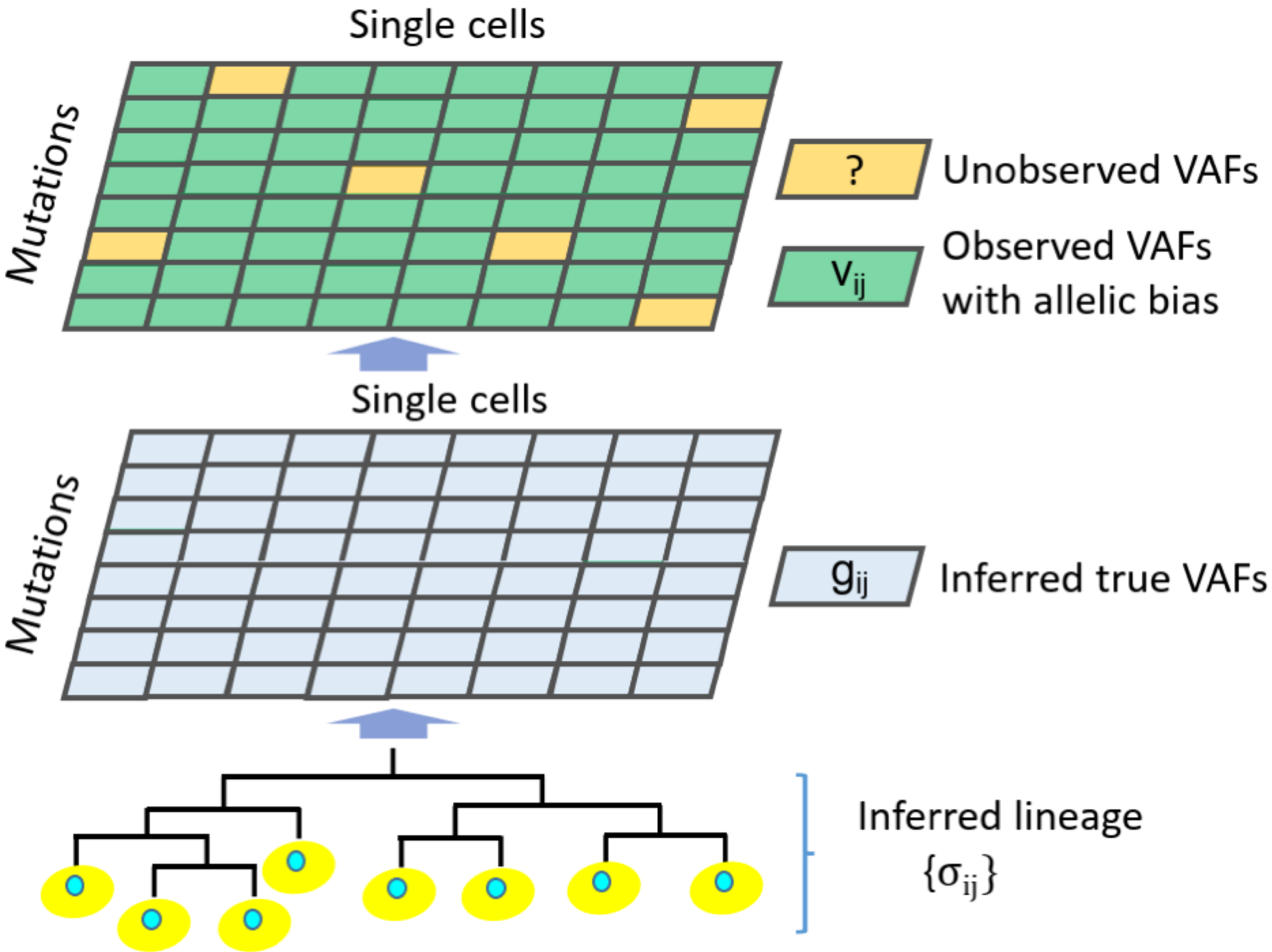

### Supplemental Figure 2

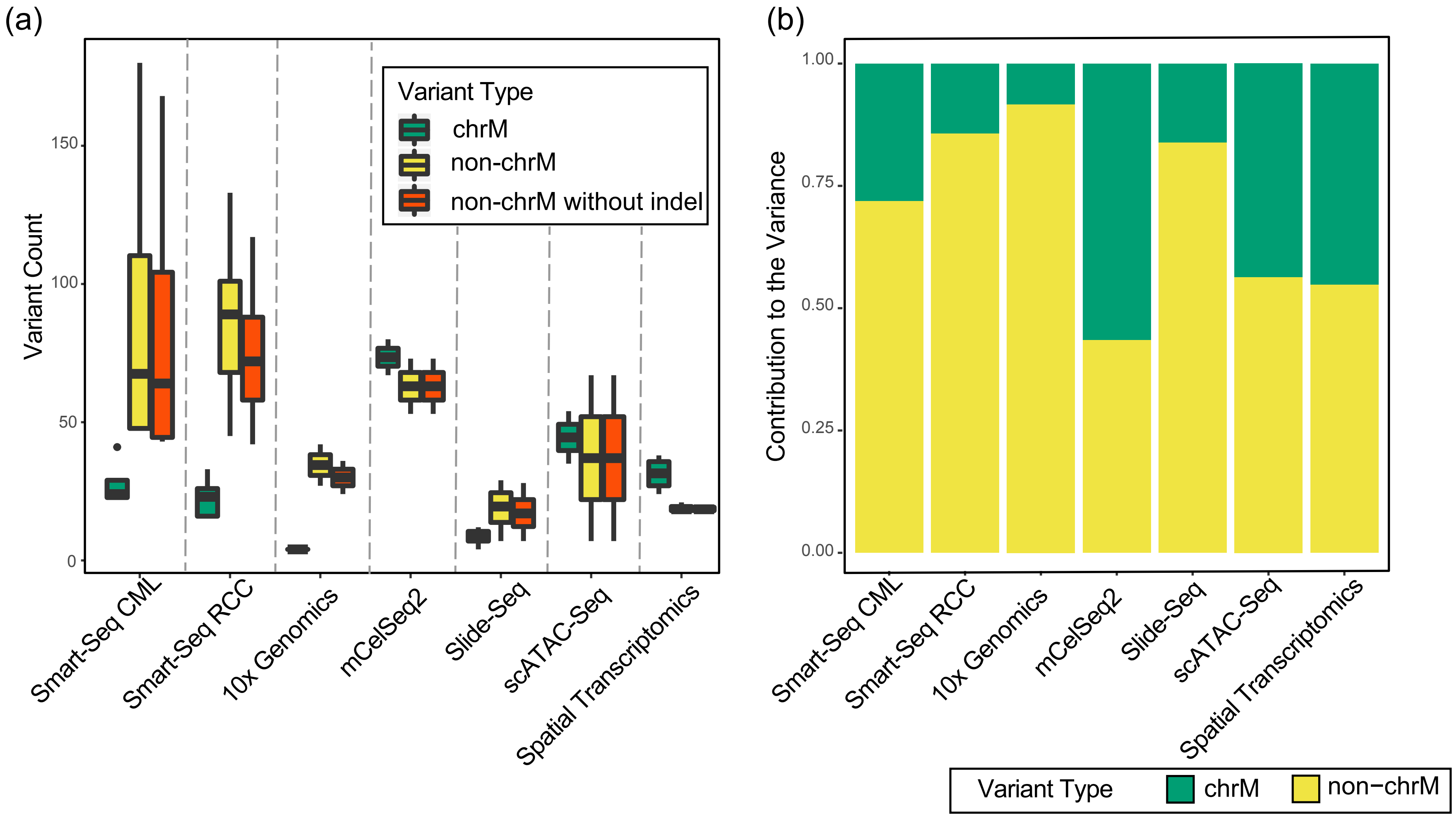

### Supplemental Figure 3

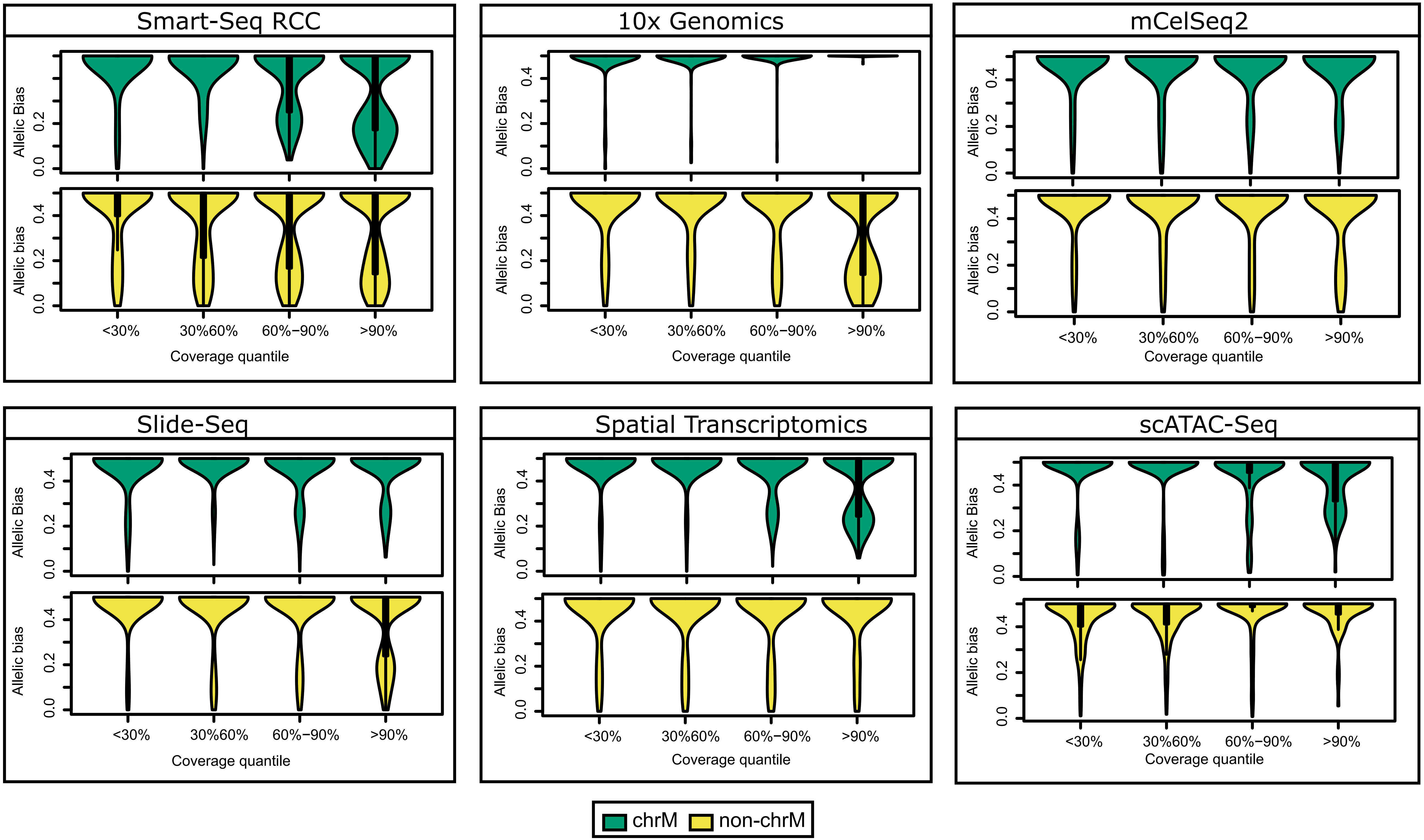

### Supplemental Figure 4

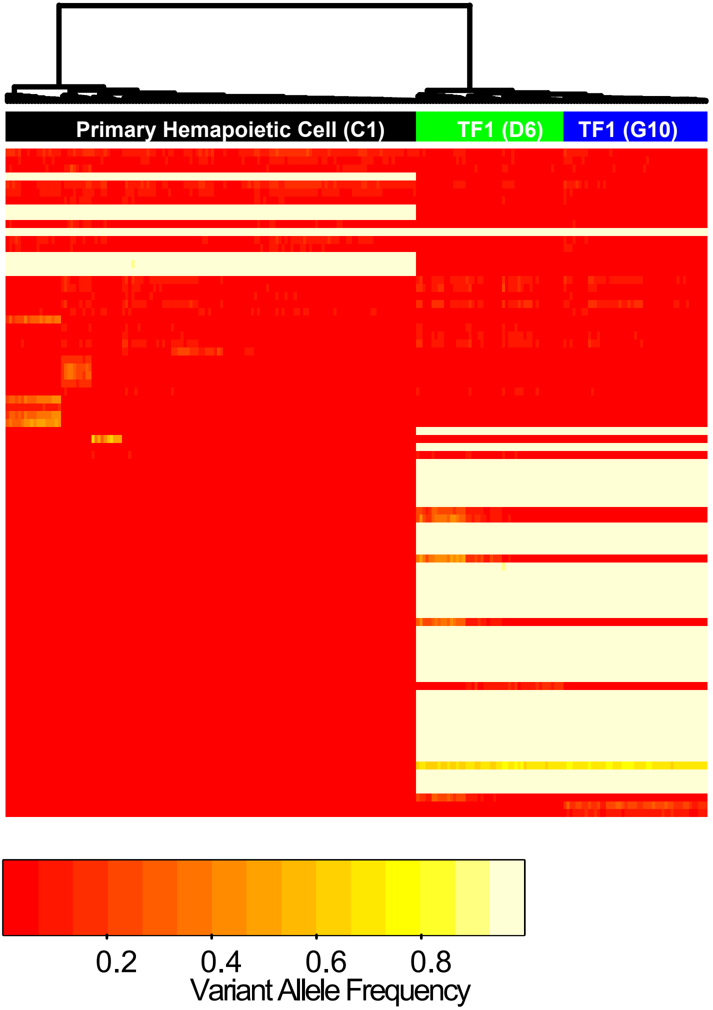

### Supplemental Figure 5

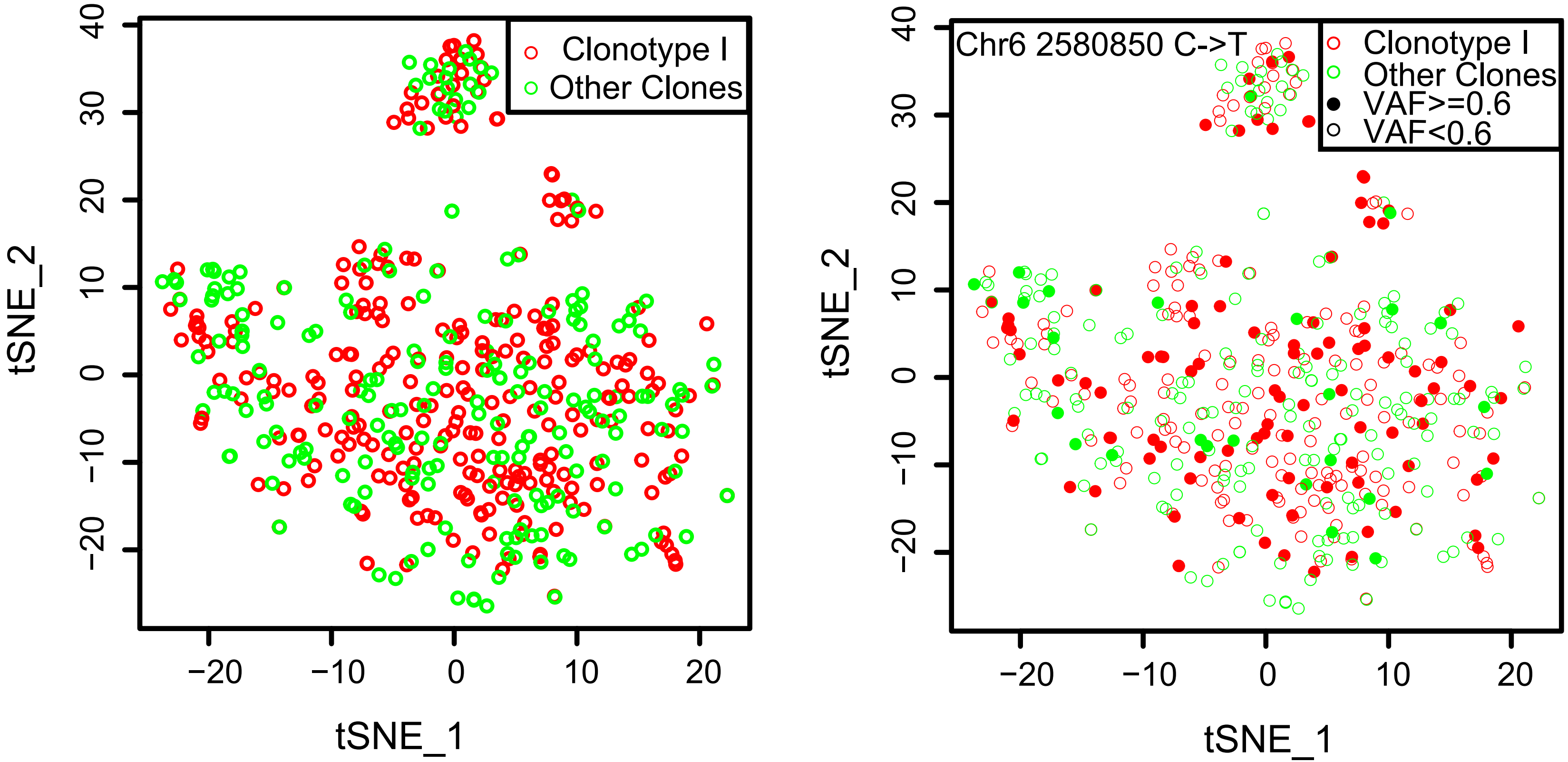

### Supplemental Figure 6

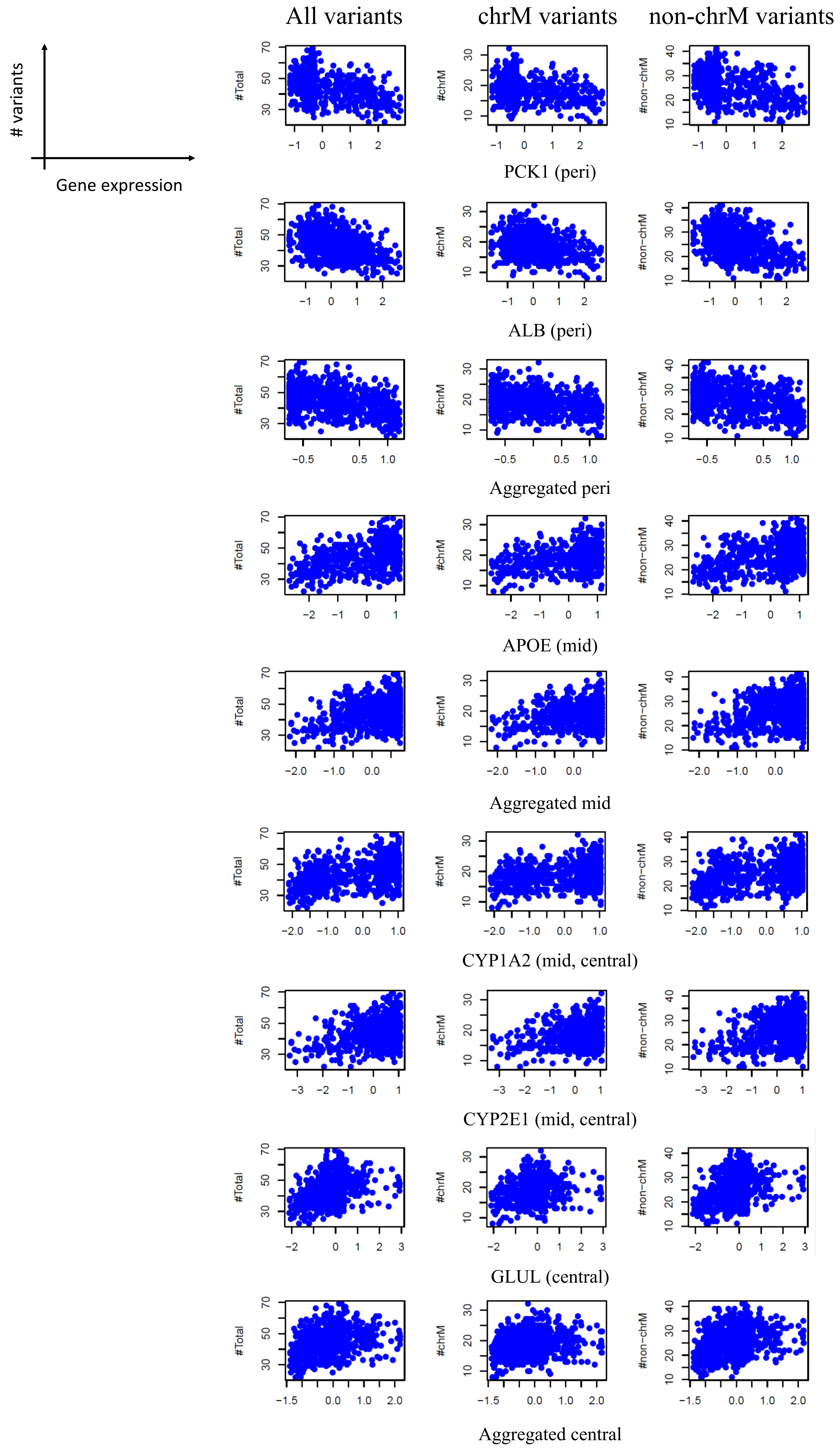

### Supplemental Figure 7

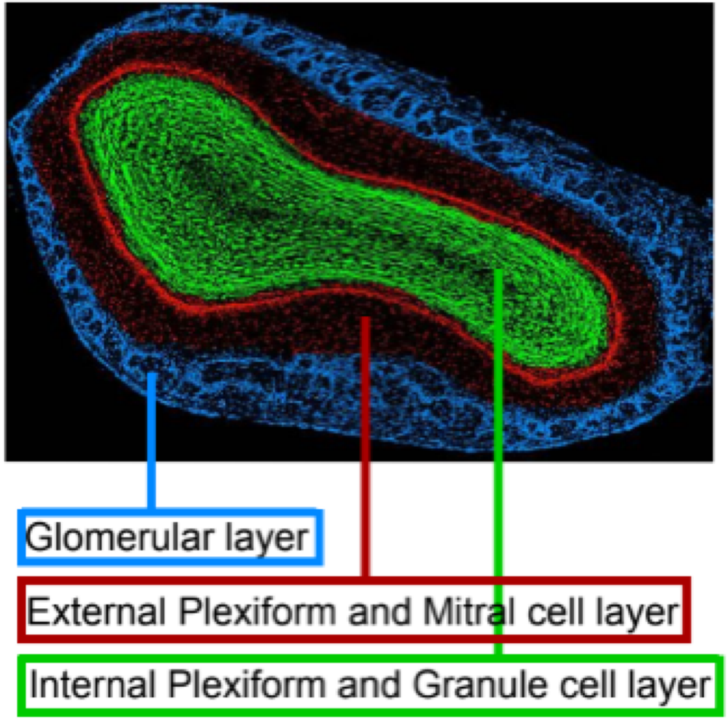

### Supplemental Figure 8

Sup. Fig. 8

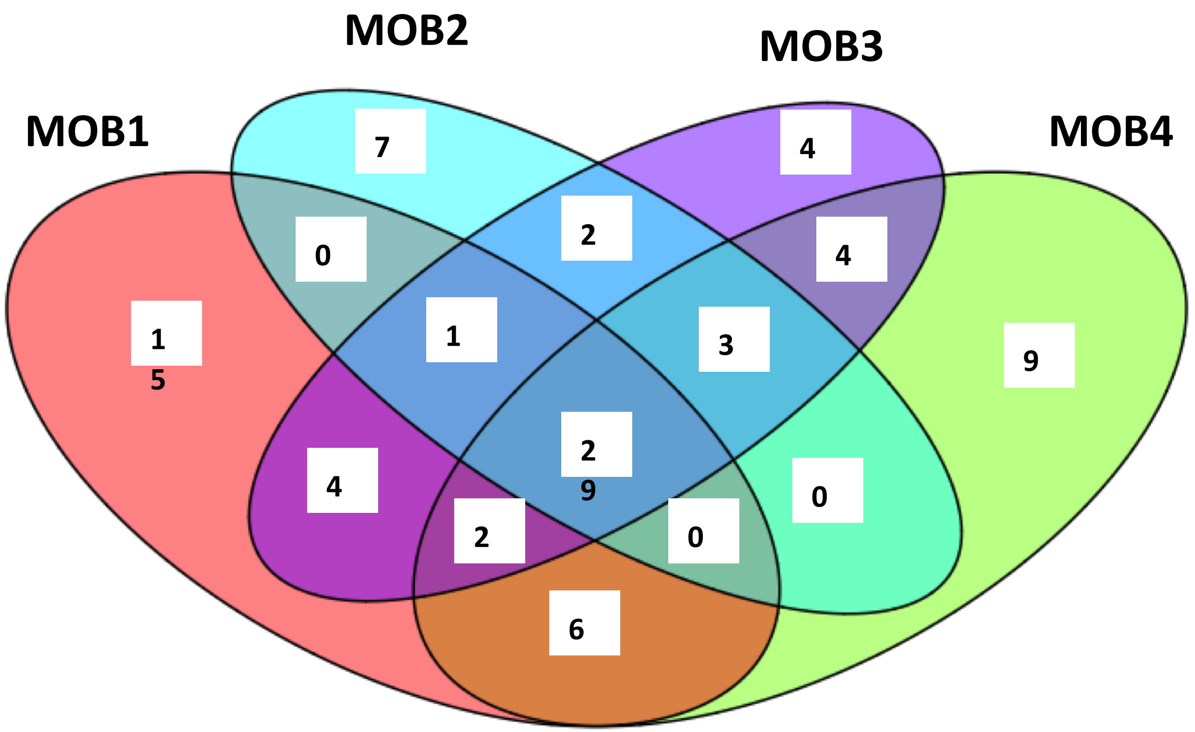

### Supplemental Figure 9

Sup. Fig. 9

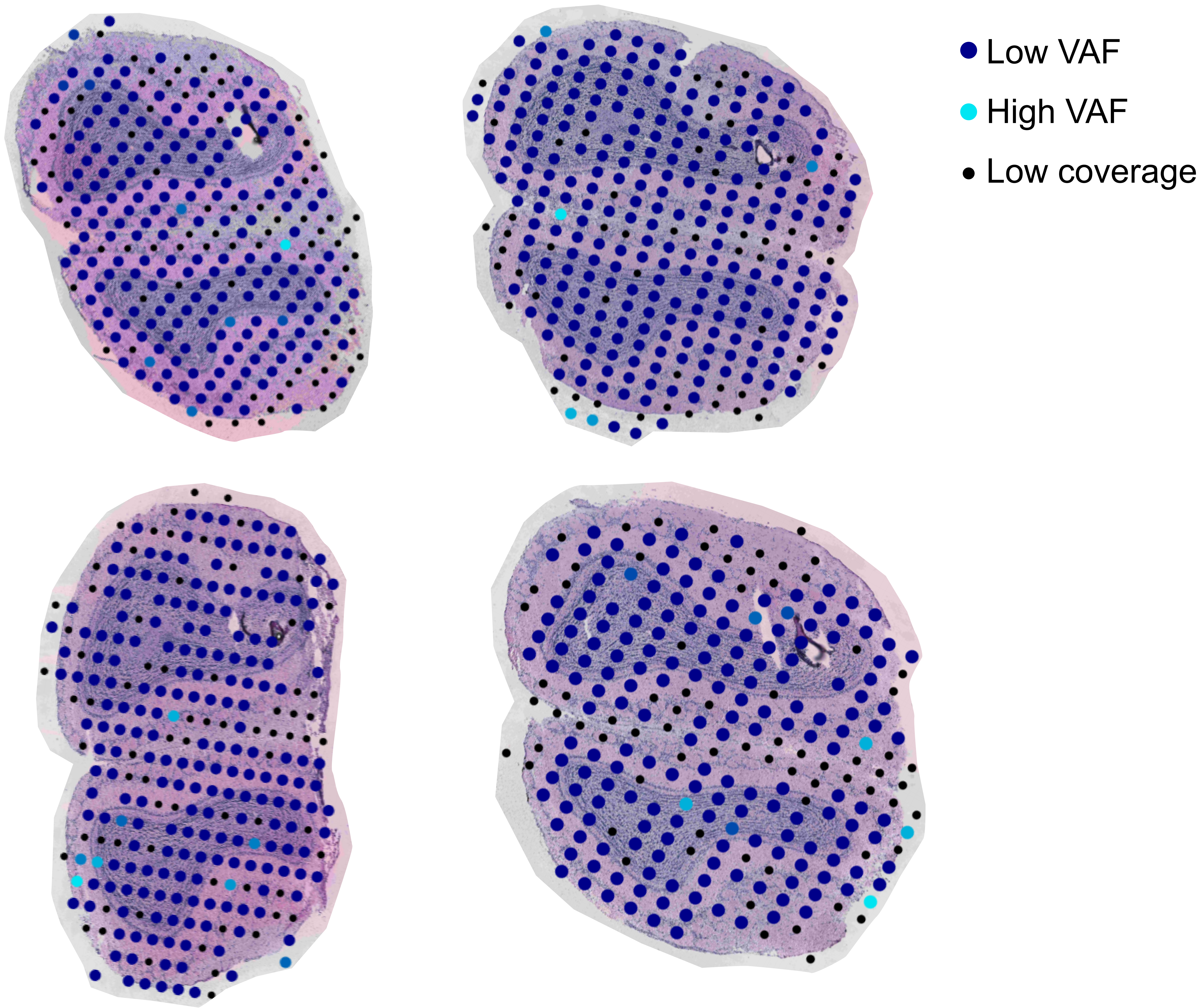

### Supplemental Figure 10

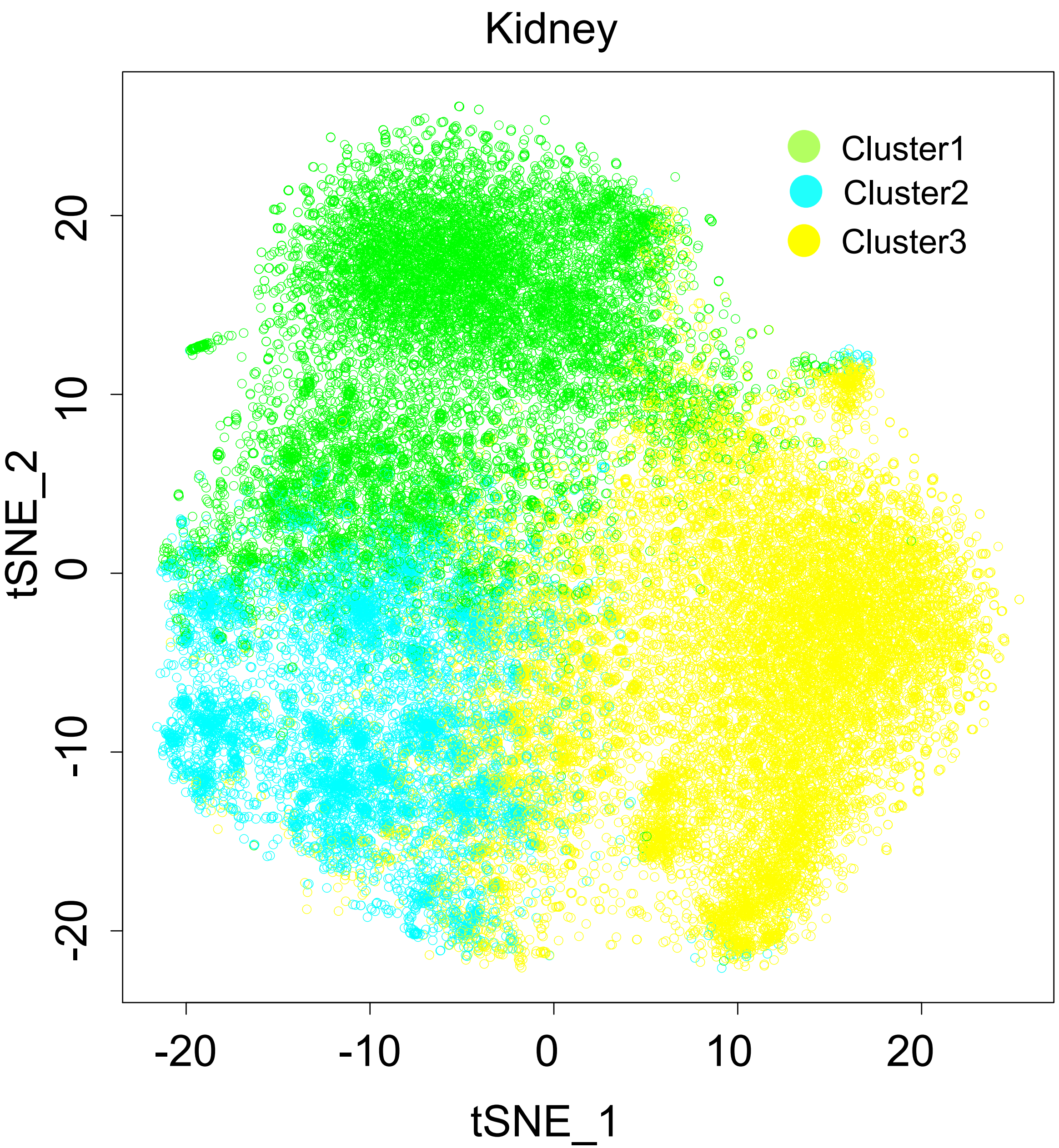

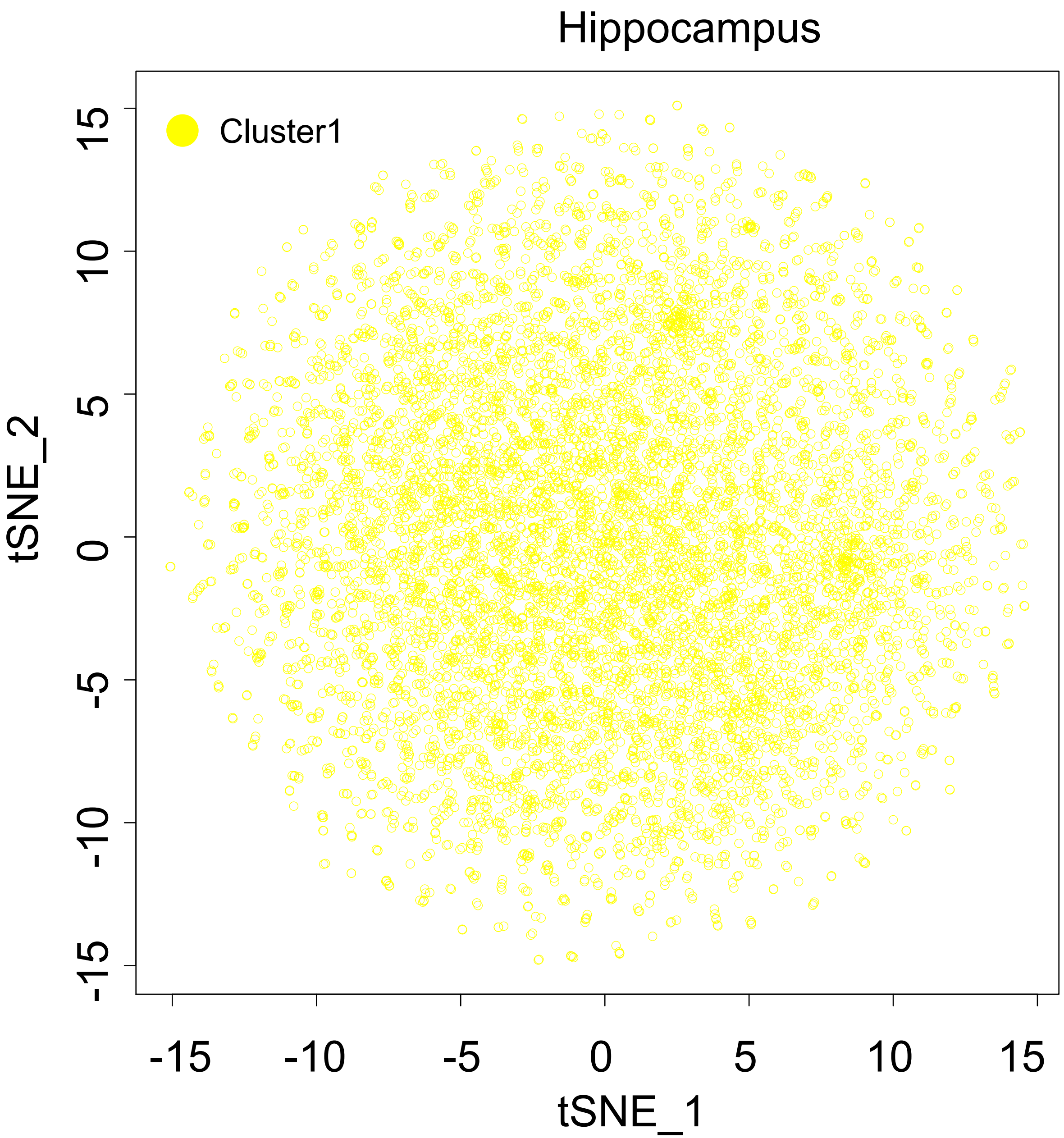

### Supplemental Figure 11

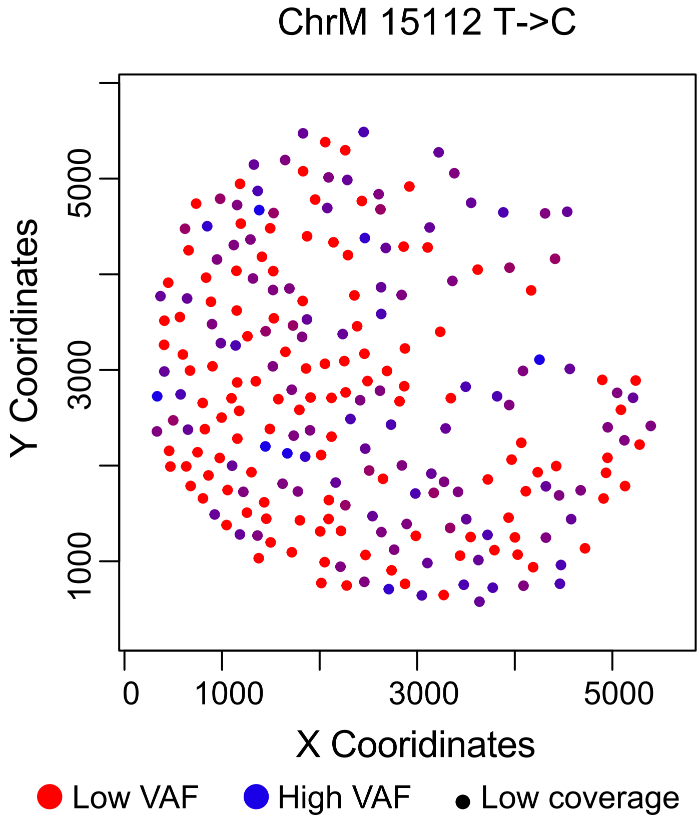
