## Supplemental File 1 for "Overcoming Genetic Drop-outs in Variants-based Lineage Tracing from Single-cell RNA Sequencing Data"

### Supplementary Material: description of the SClineager model

December 27, 2019

#### 1 Overview

Lineage tracing provides key insights into the fate of individual cells in complicated organisms. This is critical for our understanding of normal developmental processes and also diseases (with applications in glomerulosclerosis, cardiac fibrosis, cancer, etc.). Compared with prior approaches such as genetic engineering, single cell sequencing-based lineage tracing is noninvasive and provides rich information such as expression of genes for all cells at the same time. However, several significant challenges exist. First, lineage tracing based on transcriptomics is insufficient to capture the developmental paths of the mixture of cells, as cells born at different times may share similar expressional patterns. Thus, lineage tracing of cells based on genetic information is needed. Secondly, due to the low coverage of the sequencing data, detection of genetic variations for single cells is challenging. The true signal must be extracted by a robust algorithm from the noisy data. Single-cell variant calling has two inherent issues: (1) low coverage on many positions and (2) allelic bias due to true imbalanced allelic expression or sampling issues. We developed the hierarchical Bayesian model, SClineager, to perform lineage tracing of single cells based on genetics markers. This algorithm infers the true genotypes of single cell variants and also the genetic trajectories of the cells with a mechanism to correct these these two issues, which we term "genetic drop-out".

#### 2 The SClineager model

##### 2.1 Designation and input data generation

Let $i = 1, \ldots, I$ denote the cells and $j = 1, \ldots, J$ denote the variants found in all cells. Let $C = \{c_{ij} : 1 \leq i \leq I, 1 \leq j \leq J\}$ represent the total sequencing coverage on each base. When $c_{ij}$ is larger than $c$, we consider position $j$ in cell $i$ to have sufficient coverage, and the variant position is observable. $c$ is a user defined value. Let $V = \{v_{ij} | c_{ij} > c\}$ represent the variant allele frequency (VAF) matrix on observable positions. The VAF is calculated as the number of variant (non-reference genome) reads divided by the total number of reads. These are real values between 0 and 1. As the VAF data range between 0 and 1, we transformed them so the transformed VAFs can be modeled by a normal distribution. Let $T = \{t_{ij} | t_{ij} = \text{logit}(\min(\max(v_{ij}, a), 1-a)), v_{ij} \in V\}$ represent the transformed VAF matrix, where $a$ is an arbitrarily small number, like 0.01. By doing this, we have $t_{ij}$ bounded in $[\text{logit}(a), \text{logit}(1-a)]$ so that logit can be performed and extremely large positive or negative numbers can be avoided.

#### 2.2 Overall structure

We assume $\{g_{ij} : 1 \leq i \leq I, 1 \leq j \leq J\}$ represent the true genotypes to be inferred, while $t_{ij}$ deviates from the true genotype by some distance due to allelic expression bias.

For an observable position $(i, j)$ such that $c_{ij} > c$, we assume $t_{ij} | g_{ij}, c_{ij} \sim N(g_{ij}, k(c_{ij}))$, where $k(c_{ij})$ relates the possible degree of allelic bias to the sequencing coverage. As a result, SClineager implies that variants in a cell that do not have enough coverage are not useful for estimating true genotypes. $k$ can be flexibly defined, but must 1) be defined on $(0, \infty)$ and 2) be monotonically decreasing as our work has shown that positions with higher coverage tend to have less allelic bias, and 3) reflect that mutations whose observed VAFs are close to 0 or 1 (apparent homozygous wild type or mutant positions) are more likely to have allelic bias (they are actually heterozygous). For example, we used

$$k(x) = \frac{1 + 20 \cdot \text{I}(|\text{VAF} - 0.5| > 0.4)}{\sqrt{x}},$$

where the true VAF is empirically plugged-in by the observed $v_{ij}$. We write $k_{ij}$ as a shorthand for $k(c_{ij})$. The user is encouraged to try other possibilities and our R package enables this.

#### 2.3 Prior specification

We assume a symmetric positive definite matrix of $\Sigma = \big(\sigma_{ij}, 1 \leq i, j \leq I\big)$, which represents the similarity between different cells. A larger $\sigma_{ij}$ means a higher genetic similarity between two cells, or in other words, they should be more closely related on the genetic trajectory. For each $j$, the distribution of the genotypes across cells can be considered as a data generation process that conforms to the genetic similarity. Therefore, we assume $g_j = (g_{1j}, \ldots, g_{Ij})^{\text{T}} \sim N_I(\mu_j, \Sigma)$, where $N_I$ denotes $I$-dimensional multivariate normal distribution, and $\mu_j$ is a mean vector. The focus of the SClineager algorithm is to infer the similarity matrix and to recover the true genotype of cells, rather than inferring the mean genotypes across cells and variants. Therefore, we took the approach of setting each element of $\mu_j$ to be a fixed value of the average of the observed VAF values across all cells for this variant, which is a plug-in estimate of the population-level average genotype. Our analyses below show that this model setting generates parameter estimates of sufficiently good accuracy. Namely, we empirically estimate $\mu_j = \frac{\sum_{i=1}^{I} t_{ij} \text{I}(c_{ij} > c)}{\sum_{i=1}^{I} \text{I}(c_{ij} > c)} 1_I$ ($1_I = (1, \ldots, 1)^{\text{T}}$: length of $I$). It is not difficult, however, to allow uncertainty in $\mu_j$, for example, by giving a uniform prior $\mu_j \sim \text{U}(b^-, b^+)$. Here, we can set $b^{\pm} = \text{Mean}(T_j) \pm \tau \cdot \text{SD}(T_j)/|T_j|$ where $T_j = \{t_{ij} : c_{ij} > c\}$, $\tau$ can be a number like 3 or 5, and Mean($A$) and SD($A$) compute the average and standard deviation of elements in a set $A$. This approach is left as a future work and its added benefit will be tested thoroughly.

For $\Sigma$, we assume an inverse-Wishart distribution, which is a standard choice for modelling covariance matrix of normal distributions:

$$p(\Sigma|d,\Psi) = \mathrm{IW}(\Sigma; d, \Psi) \propto |\Sigma|^{-(d+I+1)/2} e^{-\frac{1}{2}\mathrm{tr}(\Psi\Sigma^{-1})},$$

where $\Psi$ is an $I$-dimensional positive definite matrix and $d$ is the degrees of freedom such that $d > I - 1$. Hyperparameters are set by $d = I$ to satisfy $d > I - 1$, and $\Psi = \mathrm{I}_I$ (an $I \times I$ identity matrix).

#### 2.4 Posterior computation

We use MCMC sampling (typically 1,000 iterations) to draw random samples from the joint posterior distribution of $G = \{g_j : 1 \leq j \leq J\}$ and $\Sigma$. The first half of the MCMC iterations are discarded as burn-in.

$$p(G, \Sigma|T, C, d, \Psi, \mu) \propto \prod_{i,j|c_{ij}>c} N(t_{ij}; g_{ij}, k_{ij}) \cdot \prod_j N_I(g_j; \mu_j, \Sigma) \cdot \mathrm{IW}(\Sigma; d, \Psi). \qquad (1)$$

Up to difference of a constant, this can be rewritten as

$$\begin{aligned} &\log(p(G, \Sigma|T, C, d, \Psi, \mu)) \\ &= -\frac{1}{2} \sum_{i,j|c_{ij}>c} \left\{ \log(k_{ij}) + \frac{(t_{ij} - g_{ij})^2}{k_{ij}} \right\} - \frac{1}{2} \sum_j \left\{ \log|\Sigma| + (g_j - \mu_j)^{\mathrm{T}} \Sigma^{-1} (g_j - \mu_j) \right\} \\ &\quad - \frac{d+I+1}{2} \log|\Sigma| \end{aligned}$$

(1) For updating $g_j$, up to a difference of a constant

$$\log(p(g_j|G_{/j}, \Sigma, T, C, d, \Psi, \mu)) = -\frac{1}{2}(g_j - \mu_j)^{\mathrm{T}} \Sigma^{-1} (g_j - \mu_j) - \frac{1}{2}(g_j - s_j)^{\mathrm{T}} K_j^{-1} (g_j - s_j),$$

where $G_{/j} = G/\{g_j\}$, $K_j^{-1}$ is a diagonal matrix with its entry being $k_{ij}^{-1}\mathrm{I}(c_{ij} > c)$ and $s_j$ is an $I$-dimensional vector with its $i$-th entry being $t_{ij}\mathrm{I}(c_{ij} > c)$. Therefore,

$$g_j|G_{/j}, \Sigma, T, C, d, \Psi, \mu \sim N_I\big(g_j; (\Sigma^{-1} + K_j^{-1})^{-1}(\Sigma^{-1}\mu_j + K_j^{-1}s_j), (\Sigma^{-1} + K_j^{-1})^{-1}\big).$$

(2) For updating $\Sigma$, up to a difference of a constant

$$\log(p(\Sigma|G, T, C, d, \Psi, \mu)) = -\frac{d+I+J+1}{2} \log|\Sigma| - \frac{1}{2}\mathrm{tr}\Big((\Psi + \sum_j (g_j - \mu_j)(g_j - \mu_j)^{\mathrm{T}})\Sigma^{-1}\Big).$$

Thus, $\Sigma$ given $G, T, C, d, \Psi, \mu$ follows an inverse-Wishart distribution:

$$p(\Sigma|G, T, C, d, \Psi, \mu) \propto \mathrm{IW}\Big(\Sigma; d + J, \Psi + \sum_j (g_j - \mu_j)(g_j - \mu_j)^{\mathrm{T}}\Big).$$

### 3 Simulation study

We conduct a simulation study to verify finite-sample performance of SClineager under various settings, which are described below. $I$ cells are observed with $J$ variants for each cell. We randomly assign $G$ biological lineages $L_g$, $g = 1, \ldots, G$, to $I$ samples. Cells in the same lineage have the same genotypes at all variant positions. Every variant is randomly assigned to have one of the three possible genotypes (assuming diploid genome) with probability 1/4 ("AA"), 1/2 ("AB"), and 1/4 ("BB"). Total sequencing coverage $c_{ij}$ is chosen in $\{1, \ldots, 40\}$, a range that is comparable to the real data VAF $v_{ij}$ is generated according to a genotype of each mutation. VAF with a genotype "AA" is from an uniform distribution $\mathrm{U}(0, 0.01)$, while that with a genotype "BB" is from $\mathrm{U}(0.99, 1)$. If a genotype is "AB", then its coverage is tested if

$$c_{ij}/40 < \mathrm{U}(0, u), \;\; u > 0; \tag{2}$$

if true (i.e., allelic bias happened), $v_{ij} = 0$ or 1 with probability 1/2, and $v_{ij} \sim N(0.5, (0.2/z_{0.005})^2)$ otherwise. The variance in the normal distribution is set to satisfy $\mathrm{P}(0.3 < v_{ij} < 0.7) = 0.99$. We record if a cell is subject to allelic bias by an indicator $b_{ij} = 0$ (no bias) or 1 (bias happened). Therefore, the allelic bias is only allowed in the genotype "AB", and it occurs more in positions with lower coverage. Note that large value of $u$ increases the bias in VAF. If the coverage is less than a pre-specified value $c$, then the corresponding VAF is unobserved and replaced by NA.

We set $G = 4$ and the proportions of lineage groups by (0.5, 0.3, 0.03, 0.17). Also, we set $c = 5$, which is close to what we chose for the real data. It is important to note that the smallest proportion we assumed is 0.03, and our analyses below show that even for this very small population of cells, the SClineager is capable of recovering their true lineages. Parameters that make different simulation scenarios are $I = 100, 200, 300$, $J = 70, 100, 150$, and $u = 0.6, 0.8, 1, 1.2, 1.4$. Each simulation scenario is repeated 20 times.

To demonstrate the distribution of the simulated data, $v_{ij}$ generated under the scheme described above when $J = 70$, $I = 100$, and $u = 1$ is presented in Figure 1. When bias exists for genotype "AB", the simulated VAF is very close to either 0 or 1, which is rather similar to those from "AA" or "BB".

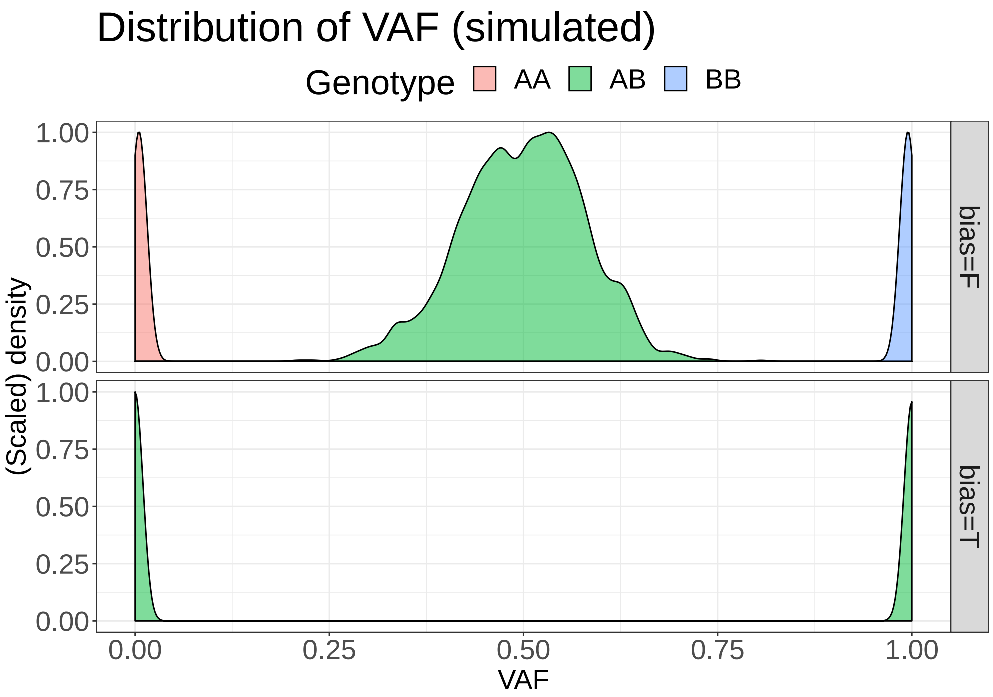


Figure 1: Density plot of the simulated VAFs for each genotype. Data is generated under $J = 70$, $I = 100$, $u = 1$. A row label "bias" indicates $b_{ij} = 0$ ("bias"=F) and $b_{ij} = 1$ ("bias"=T).

Performance of the model is evaluated for each posterior parameter. For the estimated VAF $\hat{v}_{ij}$, we compare it with the truth. The estimated $\hat{\Sigma}$ is used in an un-supervised clustering algorithm to divide samples to different lineages, which can be compared to the true lineage relationships. Specifically, we use hierarchical clustering based on a mean agglomeration method (i.e., distance between two clusters is an average of all-paired distances) where the distance of cell $i$ and $j$ is measured by $1 - \hat{\sigma}_{ij}$. In the algorithm, we fix the number of clusters to be inferred by the true value 4 (the number of true lineages). However, it is evident from a heatmap below of the estimated correlation matrix that the SClineager detected four distinct clusters of cells.

#### 3.1 Validation of posterior estimates

Figure 2 shows the fitted values of VAFs in one of the simulation scenarios ($J = 70$, $I = 100$, and $u = 1$). It is important to see that VAFs of cells and variants with allelic bias are retrieved (bottom part of each sub-panel) and their distributions are similar to those of heterozygous variants in cells with no allelic bias (top part of each sub-panel).

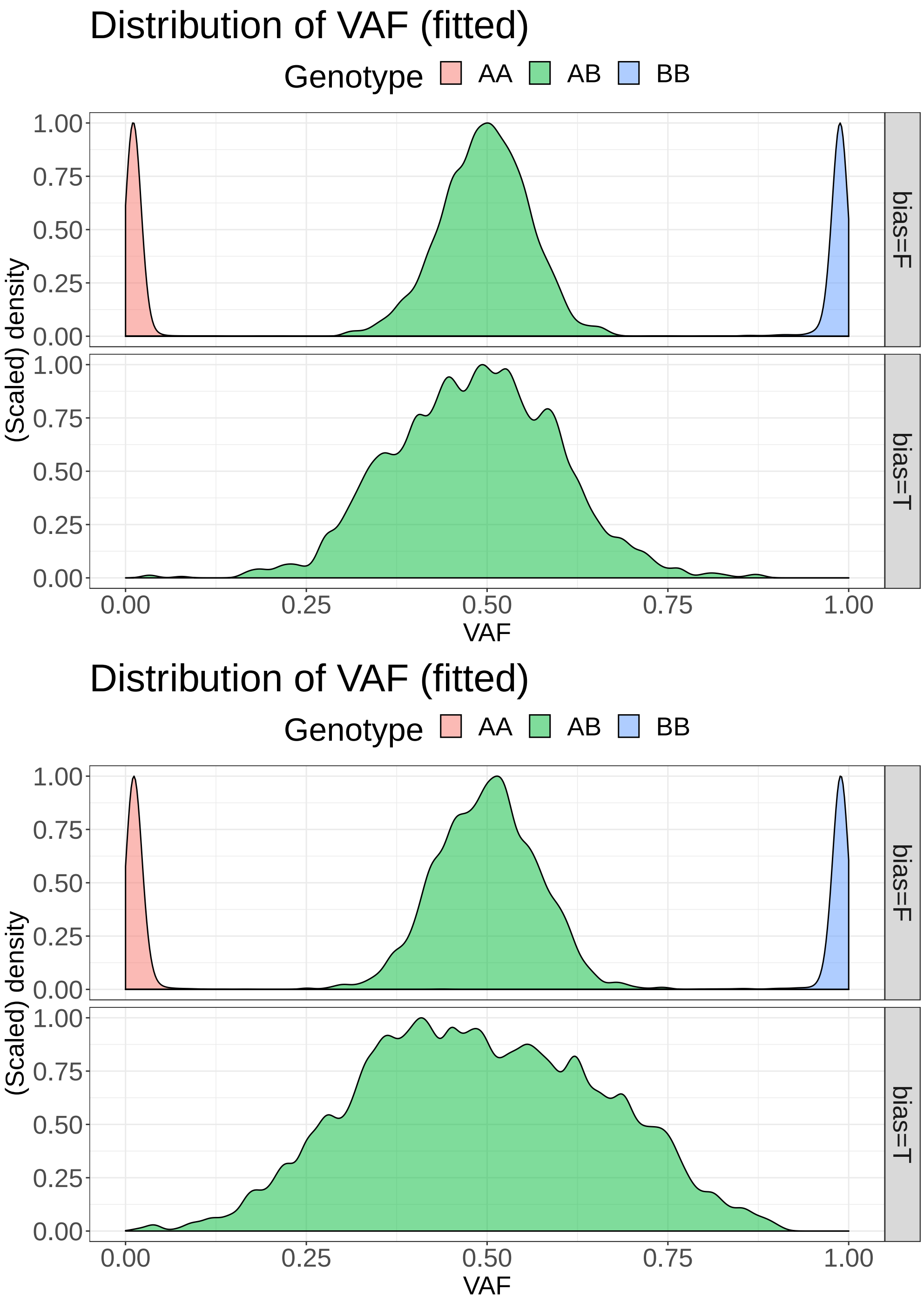


Figure 2: Density plot of the fitted (estimated) VAFs for each genotype when $u = 1$ (top) or $u = 1.4$ (bottom). Data is generated under $J = 70$, $I = 100$. A row label "bias" indicates $b_{ij} = 0$ ("bias"=F) and $b_{ij} = 1$ ("bias"=T).

We evaluated the deviance of the estimated VAFs from the true ones. Let $N_{AA} = \Big\{(i, j) :$

the genotype of $(i, j)$-th cell is "AA"$\}$. Similarly, $N_{AB}$ and $N_{BB}$ are defined. Figure 3 shows the median squared error $\text{median}_{i,j:(i,j)\in N_x}|v_{ij} - \hat{v}_{ij}|^2$ per each genotype, where $N_x$ would be switched over $N_{AA}, N_{AB}, N_{BB}$. As the results are similar for simulations created with different $J$, only the $J = 70$ case is shown in Figure 3. It can be seen that the differences between the true VAFs and the fitted VAFs are minimum for homozygous wild type ("AA") or mutant ("BB") positions. For heterozygous ("AB") positions, the differences are also very small. As extent of allelic bias increases ($u$ increases), estimation accuracy degrades for all genotypes. However, even for very large $u$, SClineager still performs reasonably well.

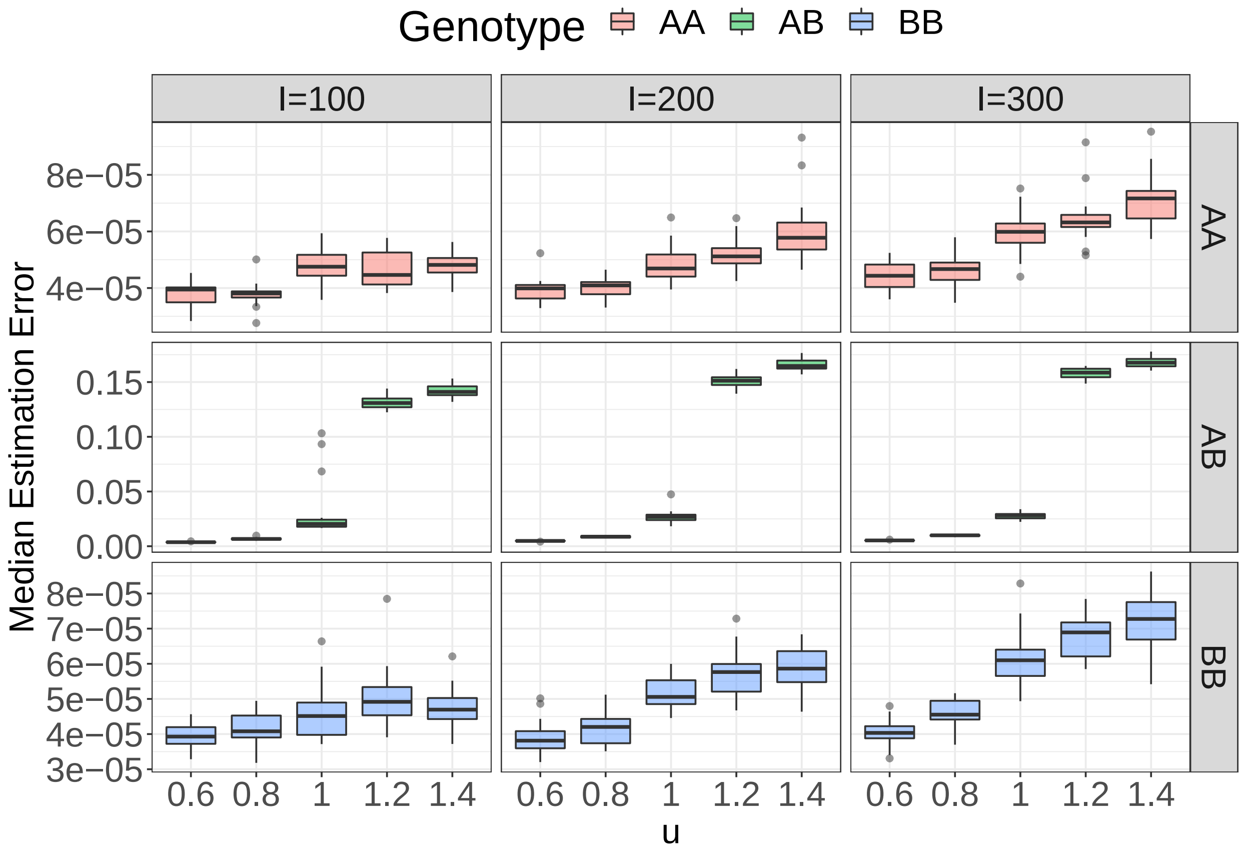


Figure 3: Median squared error of the VAFs estimated by the SClineage model when $J = 70$. The error is computed over $N_{AA}$ (top), $N_{AB}$ (middle), and $N_{BB}$ (bottom).

The accuracy of the inferred lineage labels is summarized by the adjusted Rand Index (ARI) (Hubert and Arabie 1985). If ARI computed between the true group labels and the estimated labels is close to 1, then the two lists of labels are similar to each other. Throughout all scenarios under consideration, we obtain almost perfect ARI scores (i.e., ARI $> 0.99$ with sd $< 0.01$). It is noteworthy that these high scores are not attributed to the clustering algorithm, but rather to the accurate estimation from the SClineager model, as different clustering methods lead to very similar results (data not shown). As another way of validation, Figure

4 displays a heatmap of the correlation matrix calculated from $\Sigma_0$ and $\hat{\Sigma}$, where $\Sigma_0$ is the sample covariance matrix that is calculated using the true VAFs, $\{t_{ij} : b_{ij} = 0\}$ and $\hat{\Sigma}$ is the posterior estimate of genetic similarity. Four blocks (1∼50, 51∼80, 81∼83, 84∼100) are clearly observed in the $\hat{\Sigma}$ matrix, which correspond exactly to the true lineage groups $L_1, L_2, L_3, L_4$ in $\Sigma_0$. This is another proof that the accuracy of the inferred lineage labels is independent of the clustering algorithm employed.

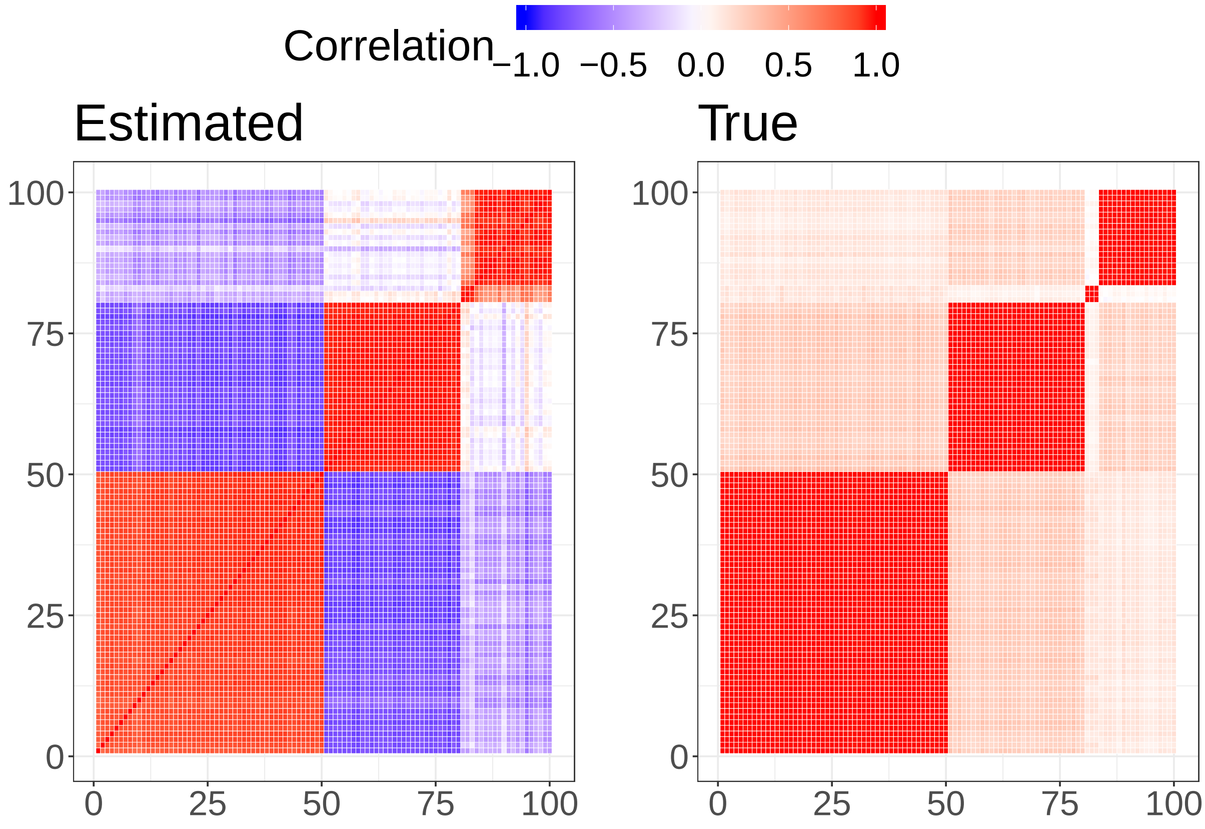


Figure 4: Heatmap of correlation matrices (100-by-100) (from $\Sigma_0$ and $\hat{\Sigma}$) of VAFs when $J = 70$, $I = 100$, and $u = 1$. The true lineage relationship can be inferred in the right matrix (red indicates the same lineage).

#### 3.2 Model diagnosis

We check the convergence of the MCMC procedure used in the SClineager model by running multiple chains with different initial starting points. We compute the posterior likelihood (1) evaluated at $\hat{G}^{(r)}$ and $\hat{\Sigma}^{(r)}$ obtained at each iteration $r$. Figure 5 shows that 8 parallel chains converge to a similar convergent point within 2000 iterations, indicating a good convergence of our MCMC algorithm.

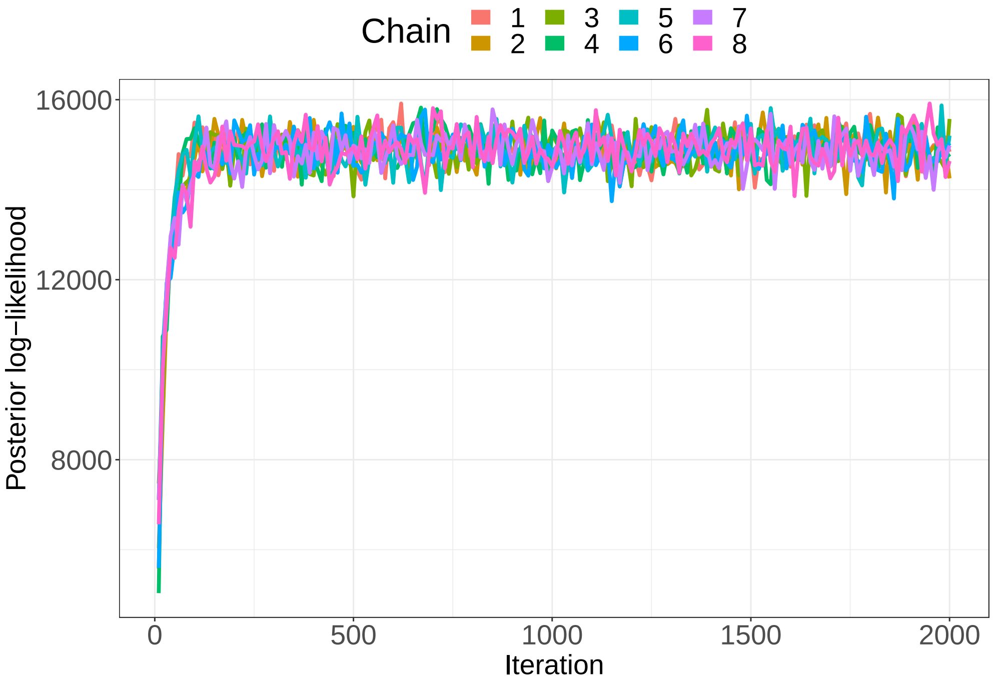


Figure 5: Multiple sequences of posterior likelihood. 8 chains of length 2000, generated with different initial conditions, are drawn for the simulation setting of $J = 70$, $I = 100$, and $u = 1$. Burn-in steps (the first 1000 iterations) are also included in the chain.

We check the goodness-of-fit using posterior predictive checking, where the key idea is to generate predictions for the parameters of the observed data points using the fitted model and test if the predictions deviate from the observed values of the same parameters. To be more specific, we use the following tail probability to measure lack-of-fit;

$$p_{ij} = \mathrm{P}\Big(t_{ij}^{new} < g_{ij} \big| T, C\Big).$$

The probability above is empirically computed by

$$\hat{p}_{ij} = M^{-1} \sum_{m=1}^{M} \mathrm{I}\big(t_{ij}^{new,(m)} < t_{ij}\big)$$

where $t_{ij}^{new,(m)}$ is generated from $N(g_{ij}^{(m)}, k_{ij})$ with $g_{ij}^{(m)}$ the $m$-th posterior sample of $g_{ij}$. Figure 6 shows the boxplot of 100 $\hat{p}_{ij}$'s that are randomly selected among cells $\{(i,j) : b_{ij} = 0\}$. The figure shows that none of these probabilities is less than 0.05 or greater than 0.95, indicating that the fitted SClineager model generates parameter predictions that are consistent with the observations. Therefore, we conclude that there is no lack-of-fit.

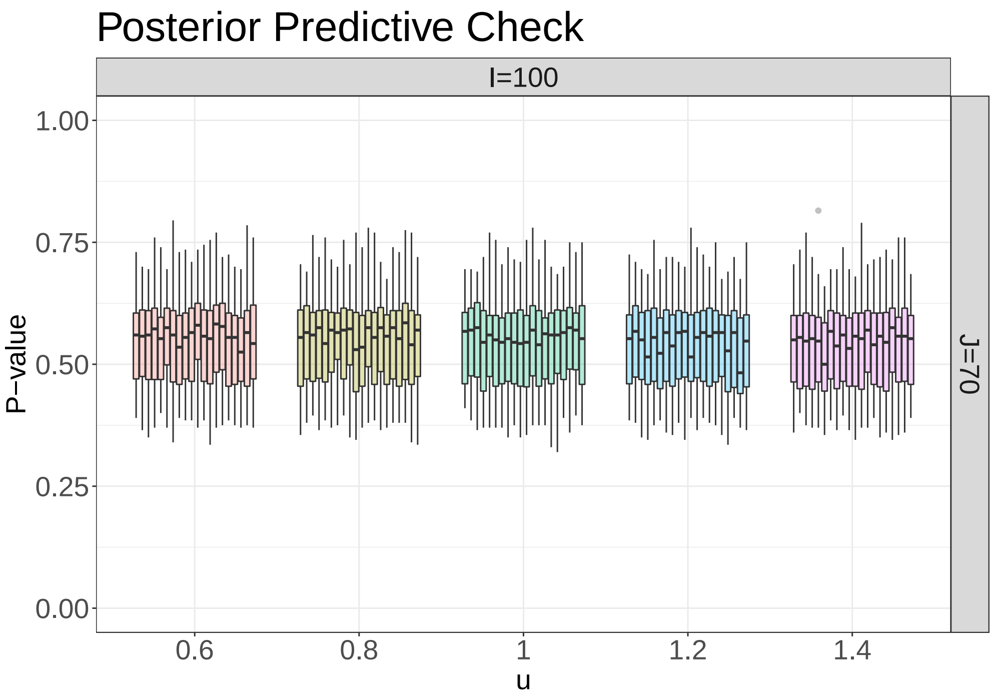


Figure 6: Boxplots of posterior predictive probabilities, which can be interpreted also as p-values. 100 randomly chosen cells are summarized in each boxplot. P-values are concentrated around 0.6, which indicates minimum evidence against lack-of-fitting.

#### 3.3 Sensitivity analysis for the cutoff value $c$

The paramter $c$ is chosen in a somewhat arbitrary manner. We tend to use a higher $c$ for single cell datasets with better coverage. We examine how sensitive the estimation outcomes of SClineager are according to different $c$ values. We apply different cutoff values to the same $\{v_{ij} : \forall i, j\}$ and get $V_c = \{v_{ij} : c_{ij} > c\}$ with $c = 3, 5, 7, 10$. Then, we apply SClineager on each of $V_c$'s and evaluate its performance, which is shown in Figure 3. In this experiment, we compute $Q_r\Big(\big\{|v_{ij} - \hat{v}_{ij,c}|^2 : (i, j) \in N_x, v_{ij} \in V_c\big\}\Big)$ for each genotype $x = AA, AB, BB$, where $Q_r(D)$ $(0 \leq r \leq 1)$ is the quantile function that returns the $100r\%$ percentile of $D$ and $\hat{v}_{ij,c}$ is the estimator of $v_{ij}$ based on $V_c$. We choose three values of $r = 0.25, 0.5, 0.75$, which yield $Q_1$, $Q_2$, and $Q_3$ of squared errors, respectively. Figure 7 shows that the estimation accuracy does not significantly change as $c$ increases. The robust performance of SClineager with respect to $c$ is also confirmed via the ARI scores that are all equal to 1 under different choices of $c$'s.

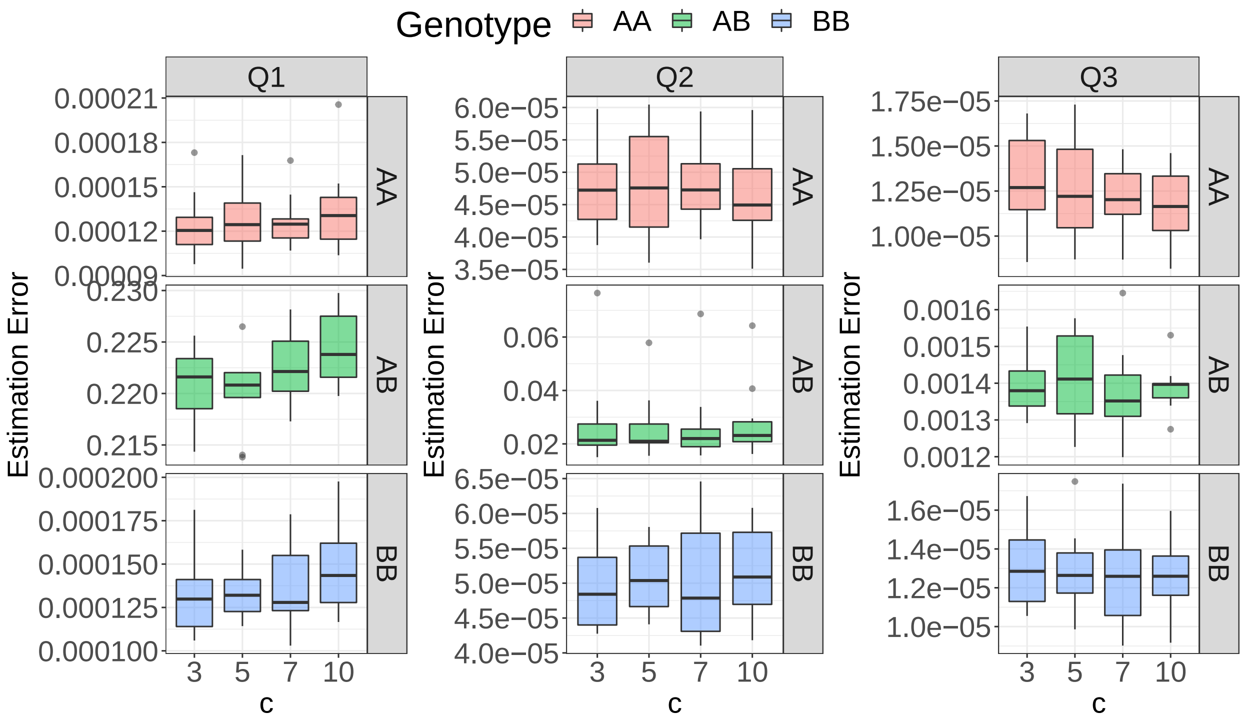


Figure 7: Quantiles of squared error of the estimated VAFs when the cutoff value is varied over $c = 3, 5, 7, 10$. $J = 70$, $I = 100$, and $u = 1$ are assumed. The error is computed over $N_{AA}$ (top), $N_{AB}$ (middle), and $N_{BB}$ (bottom).

#### 3.4 More complicated scenarios

The simulation of four mutually exclusive lineages may be somewhat simplistic compared with the true lineages that can be expected in real data, where one lineage of cells can be further split into more than one sub-lineages. These sub-lineages are different but still similar to each other, and together, they are different from the other lineages (which can also have their own sub-lineages). To reflect this, we further consider sub-structure in lineages by assuming a split within a lineage into two of equal size. The genotypes of these two sub-lineages are different in the first 25% of genotype sequences, but are the same in the other 75% (see Figure 8).

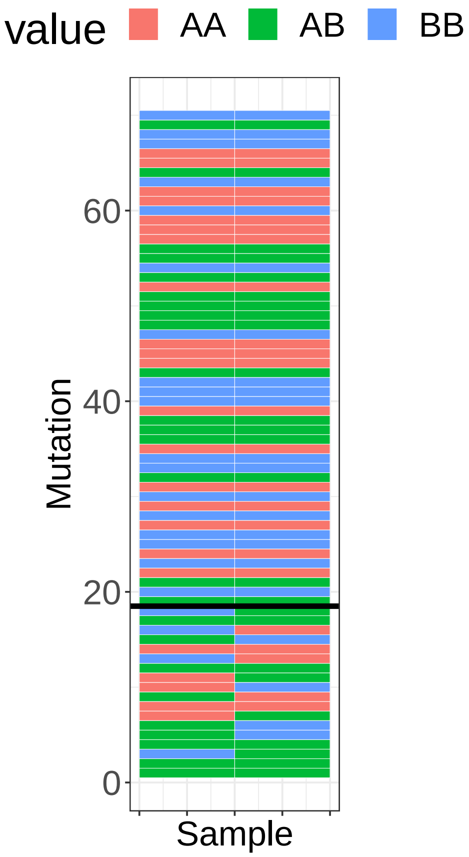


Figure 8: The genotypes of the two sub-lineages. The last 75% pairs (above the black line) of genotypes are identical since these two sub-lineages belong to the same parent lineage, while the other genotypes (below the black line) are different as these two sub-lineages will diverage from each other to some extent.

In this simulation task, we assume a complicated lineage structure with several such sub-lineage splits, which is visualized below.

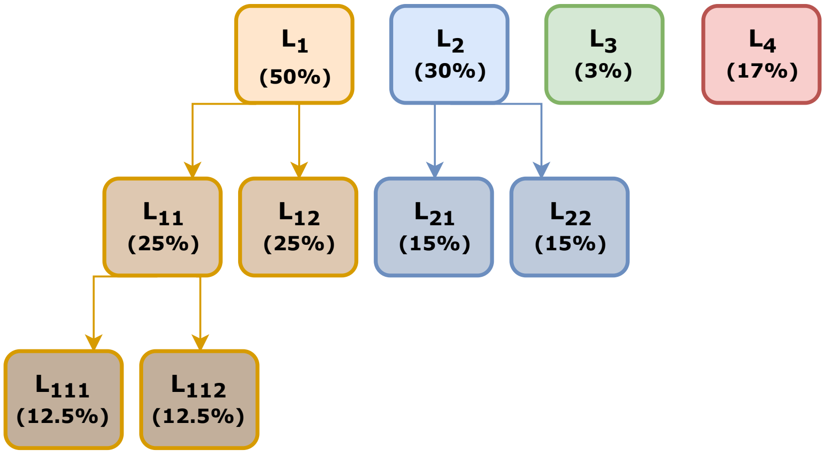


Figure 9: Tree diagram for the assumed lineage structure of the single cells. Percentage in parenthesis is the proportion of cells in each lineage.

The variant and VAF observations are generated as before, where we fix $I = 100$ and $J = 70$ and vary $u = 0.6, 0.8, 1, 1.2, 1.4$.

Figure 10 displays the true lineage structure and the estimated correlation matrix of the simulation dataset above. Four main blocks can be clearly discerned in both matrices. A closer examination at the first block (1$\sim$50) or the second (51$\sim$80) shows sub-blocks of cells, which correspond to their respective sub-lineages. We compute ARI scores to access detection of the (sub-)lineage structure. By setting the number of clusters by 4, the hierarchical clustering algorithm finds four main lineages with ARI $> 0.88$ (sd $< 0.14$) in all cases. When the cells are divided into 7 groups by the clustering algorithm, ARI achieves $> 0.90$ (sd $< 0.1$) in all cases.

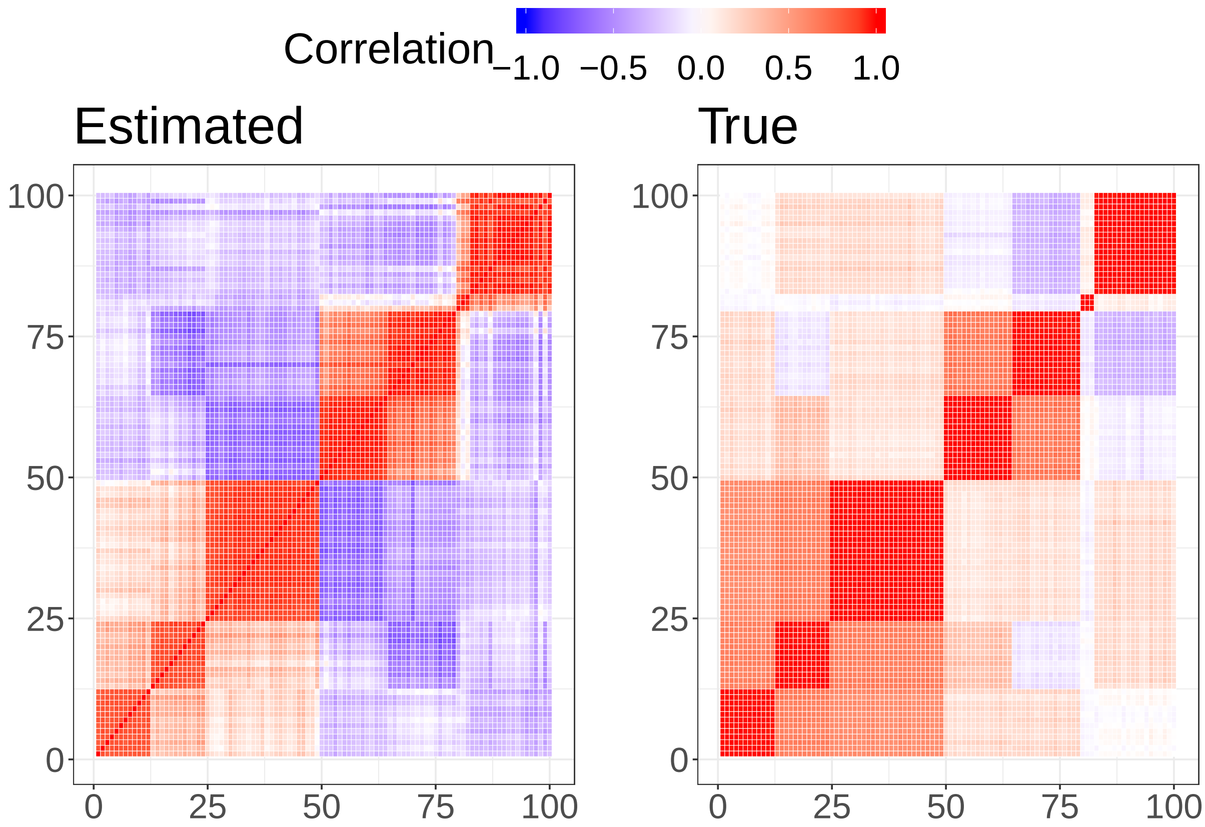


Figure 10: Heatmap of correlation matrices (100-by-100) (from $\Sigma_0$ and $\hat{\Sigma}$) of VAFs of the sub-lineage simulation task when $J = 70$, $I = 100$, and $u = 1$. The true lineage relationship can be inferred in the right matrix (red indicates the same lineage).

We also show the fitted VAFs of the variants in single cells and the convergence of posterior estimates, which shows that the accuracy of SClineager in such scenario is similar to its performance when no sub-lineage structure is assumed. (see Figure 11 and 12)

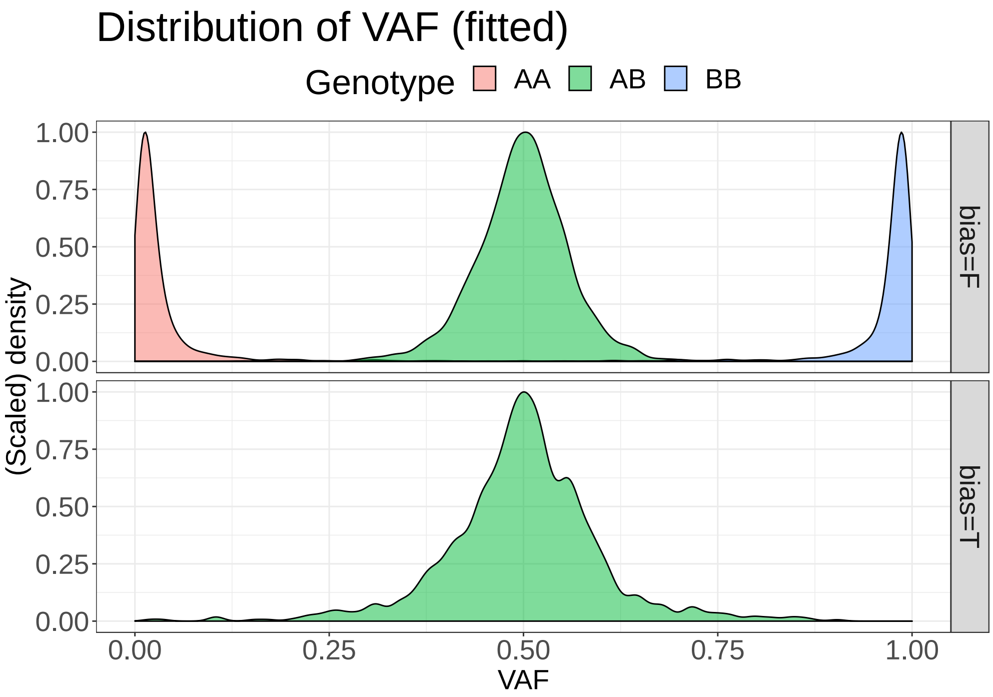


Figure 11: Density plot of the fitted VAFs for each genotype class, in the sub-lineage simulation task, where $J = 70$, $I = 100$, $u = 0.6$ is assumed. The fitted (estimated) VAF data are displayed. A row label "bias" indicates $b_{ij} = 0$ ("bias"=F) and $b_{ij} = 1$ ("bias"=T).

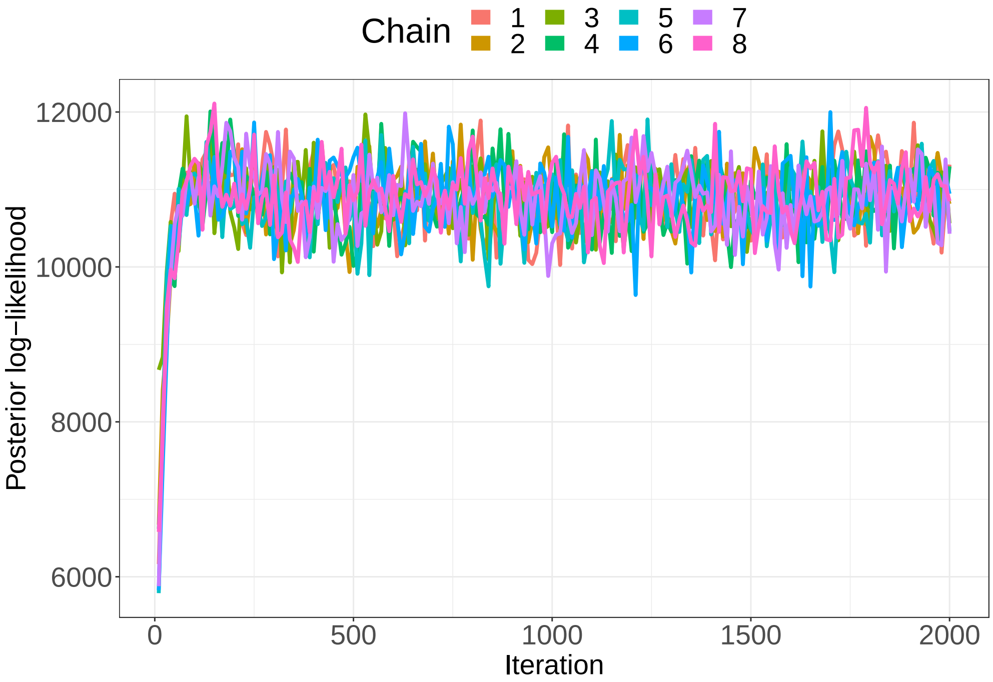


Figure 12: Multiple sequences of posterior likelihoods generated for the sub-lineage simulation task. 8 chains of length 2000, generated with different initial conditions, are drawn, for the simulation setting of $J = 70$, $I = 100$, and $u = 1$. Burn-in steps (the first 1000 iterations) are also included in the chain.

Together, these results further prove that the SClineager model can handle lineage tracing of single cells based on genetic markers in scenarios of complicated linegae structures that approximate true single cell sequencing data.
